# The host range paradox of *Meloidogyne incognita*: a physiological and transcriptomic analysis of nine susceptible interactions across six plant orders

**DOI:** 10.1101/2025.09.26.678250

**Authors:** Victor Hugo Moura de Souza, Clement Pellegrin, Vincent C.T. Hanlon, Chongjing Xia, Olaf P. Kranse, Unnati Sonawala, Priya Desikan, Beatrice Senatori, Etienne G.J. Danchin, Lida Derevnina, Sebastian Eves-van den Akker

## Abstract

The plant-parasitic nematode *Meloidogyne incognita* is the pathogen with the broadest host range among all known biotrophic interactions. This species is also the single most damaging of a group of agriculturally important plant-parasites, which together are estimated to contribute to losses in excess of $170 billion/year to world agriculture. Understanding how *M*. *incognita* is able to infect representatives from most orders of flowering plants, covering more than 3000 species, addresses a fundamentally important question of how pathogens adapt to their host-environment, and may inform control of a pathogen which threatens global food security. Here, we analyse the plant-nematode infection phenotype, and cross-kingdom transcriptome, of nine interactions across six orders of flowering plants at 25 days post infection. At this stage, majority of nematodes found within roots were immature and mature females (49.2% - 91.8%). Our data show that the phylogenetic distribution of hosts does not explain the phenotypic distribution of parasitism. Interestingly, though, *M. incognita* do have distinct transcriptional responses to different groups of hosts, but in a pattern which is independent of host phylogenetic. Three distinct nematode “transcriptional programmes” - Group 1, 2, and 3 - are evident, and we find that effectors are neither uniformly deployed across hosts, nor across groups of hosts. Importantly, we show that this differential deployment of effectors can have profound consequences for host specificity. Finally, we show that there is essentially no widespread core gall transcriptome at 25 days post infection, prompting the proposal of a model best described as “all roads lead to Rome”.

## Introduction

Of all biotrophic interactions, the pathogen with the broadest host range is the plant-parasitic nematode *Meloidogyne incognita*. *Meloidogyne incognita* is also the single most damaging of a group of agriculturally important plant-parasites, which together are estimated to contribute to losses in excess of $170 billion/year to world agriculture^1^. Understanding how *M. incognita* is able to infect representatives from most orders of flowering plants, covering more than 3,000 species^2^, may therefore inform control of a pathogen which threatens global food security, while shedding light on the fundamentally important question of how pathogens adapt to their local host-environment^3^.

It is widely accepted that infection follows a defined series of events, regardless of host. The second-stage juveniles (J2) of *M. incognita* penetrate roots, move towards the root tip, turn around, and move basipetally until they find a suitable group of cells to induce the formation of the feeding site (termed “giant cells”)^4^. Concurrent with feeding site establishment, J2s secrete a large pool of effector proteins from specialised esophageal glands, delivering them into and around host cells through a hollow, spear-like stylet^5^. A group of host cells in the parenchyma vascular tissue is re-differentiated, through repeated mitosis without cytoplasm division, into so-called giant cells. Several of these abnormally large cells each contain up to one hundred enlarged nuclei, and are embedded in hypertrophied proliferated tissue termed a gall^4^. Abundant nutrients supplied by the feeding site enable females to lay 500 - 2000 eggs, within a gelatinous matrix^6^.

Remarkably, *M. incognita* is capable of inducing this novel pseudo-organ – multiple giant cells within a gall – in each of ∼3,000 different hosts, principally in roots but in some cases even stems and leaves^7^. Antithetical to this exceptional host diversity is an apparent absence of pathogen diversity: *M. incognita* reproduces exclusively via parthenogenesis and therefore has limited ability to recombine genetic variation^8,9^. The host range paradox is therefore best described by the juxtaposition of extreme host range, highly specialised parasitism, and yet, clonal reproduction. Interestingly, *M. incognita* is a triploid species with three AAB sub-genomes, which arose due to hybridisation events^10,11^. These hybridisation events resulted in most genes being initially present as three homologous copies, yielding an expanded effector repertoire, and is hypothesized to contribute to the exceptional host range of *M*. *incognita*^11^.

Here we analyse the infection phenotype and transcriptome of nine plant-nematode susceptible interactions across six orders of flowering plants at 25 days post infection (dpi). Our data show that the phylogenetic distribution of the hosts does not explain the phenotypic distribution of parasitism. Nematodes do, however, have distinct transcriptional responses to different groups of hosts, but in a pattern which is independent of host phylogeny and phenotypic similarities. Three distinct nematode “transcriptional programmes” – Group 1, 2, and 3 – are evident, and we find that effectors are neither uniformly deployed across hosts, nor across groups of hosts. Importantly, we show that this differential deployment of effectors from the different sub-genomes can have profound consequences for host specificity. Finally, we show that there is essentially no widespread core gall transcriptome at 25 dpi, prompting the proposal of a model best described as “all roads lead to Rome”.

## Results

In an attempt to understand how *M. incognita* is able to form prolonged biotrophic interactions on thousands of host species^2^, we analysed the physiology and gene expression of host and parasite in nine susceptible interactions across six orders of flowering plants, under identical growing conditions. We recorded and analysed several phenotypic traits of *M. incognita* infection at 25 dpi: gall appearance; gall ratio (gall width/adjacent root width); gall index^12^; number of nematodes per gall; and the developmental progression of infecting nematodes (Fig.1).

**Figure 1.**
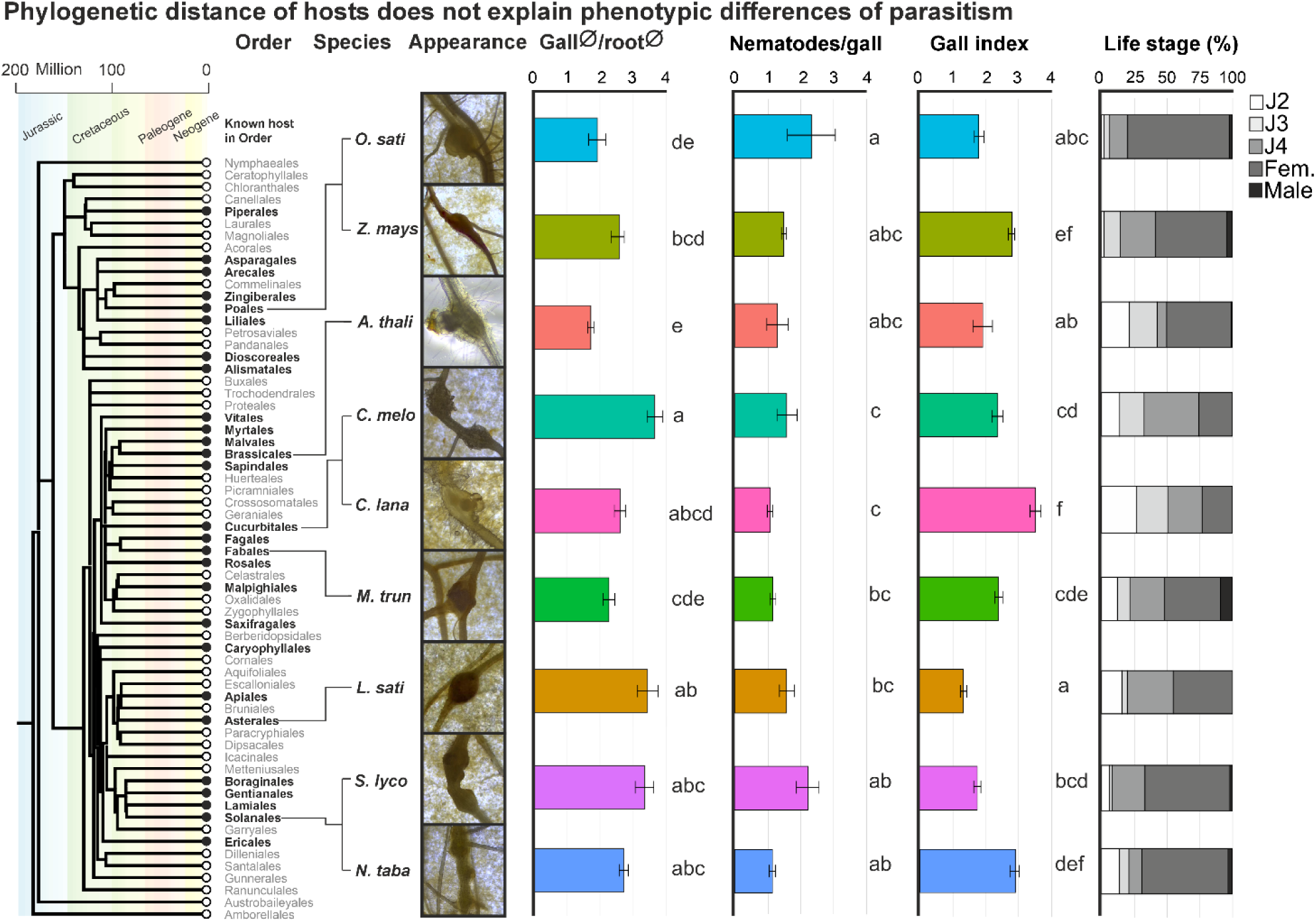
Phylogenetic distance of host does not explain phenotypic differences of *Meloidogyne incognita* parasitism at 25 dpi. Phylogenetic tree of flowering plants (https://timetree.org) highlighting *M. incognita* host distribution, in which black dots mark orders containing at least one host plant infected by *M. incognita*. Observed phenotypic differences (gall appearance, Gall ratio (Log (x+1) of gall diameter/root diameter(⌀)), nematodes per gall, gall index^12^, and life stage percentage) are shown. Error bars indicate the standard error. Differences in gall ratio were identified by ANOVA and *post-hoc* Tukey’s test. Lower case letter indicate homogeneous subsets. Differences in all other parameters were identified by Kruskal-Wallis test, followed *post-hoc* Dunn’s test.

Surprisingly, observed similarities and differences between host responses do not consistently match the phylogeny of the hosts. For example, while maize and melon diverged 163.3 Million Years Ago (MYA)^11^ they present statistically indistinguishable gall ratios, while Arabidopsis and melon diverged 121 MYA^11^ yet their gall ratios were significantly different (ANOVA *post-hoc* Tukey’s test p < 0.001). Similarly surprising, the rate of nematode development differed between plant species, but was both inconsistent with the phylogeny of those species and the afore mentioned infection phenotypes of the host. For example, even though the lowest proportion of nematodes that developed to mature females at 25 dpi was observed in the relatively closely related melon and watermelon (last common ancestor 20.4 MYA^13^), they significantly differed from rice, tobacco and tomato (Kruskal-Wallis *post-hoc* Dunn’s test p < 0.05) which share a last common ancestor 163.3 MYA^13^, and do not differ from one another. Taken together, the host’s physiological response to nematode parasitism, and the nematode’s developmental progression, are at least in part decoupled from host evolutionary history.

To explore these nematode and host differences, a cross-kingdom RNAseq experiment was performed on all nine susceptible interactions. The experiment was designed to focus on the local effects of prolonged biotrophy, where galls were collected from each host at 25 dpi, as well as adjacent equivalent uninfected segments of roots from the same plants. RNA was extracted, sequenced, and mapped to the corresponding host and nematode genomes to enable the measurement of plant gene expression, and nematode gene expression, from the same sample.

Focusing initially on nematode expression, principal component analysis (PCA) of normalized read counts revealed three “programmes” of gene expression (Fig. 2): Group 1: rice, tobacco, tomato and watermelon (Mesangiospermae LCA ∼ 160 MYA), Group 2: Arabidopsis and Medicago (Rosids LCA ∼ 108 MYA), and Group 3: Lettuce, maize and melon (Mesangiospermae LCA ∼ 160 MYA)^13^. This clustering, is surprisingly inconsistent with host phylogeny and nematode phenotypes. For example, the nematodes infecting the monocots rice and maize were separated into Groups 1 and 3 respectively, as were those infecting the Cucurbits watermelon and melon. While we explored other clustering methods, (e.g. those based on raw read counts or hierarchical clustering scores (hclust)), which provided better, but still imperfect, matches to host phylogeny (Fig. S1), the variance stabilizing and regularised log transformations (e.g. Fig. 2) were more consistent with pairwise differential gene expression between groups. For example, there are more *M. incognita* differentially expressed genes between rice and maize (different groups; 7,517) than between melon x maize (same group; 255). Given that commonly used transformations result in the clustering shown in Fig. 2, and these are consistent with the pairwise analysis of nematodes on each host, we favour this analysis, but present alternatives (Fig. S1) as further evidence for inconsistency of host phylogenetic signal in nematode transcriptional programmes.

**Figure 2.**
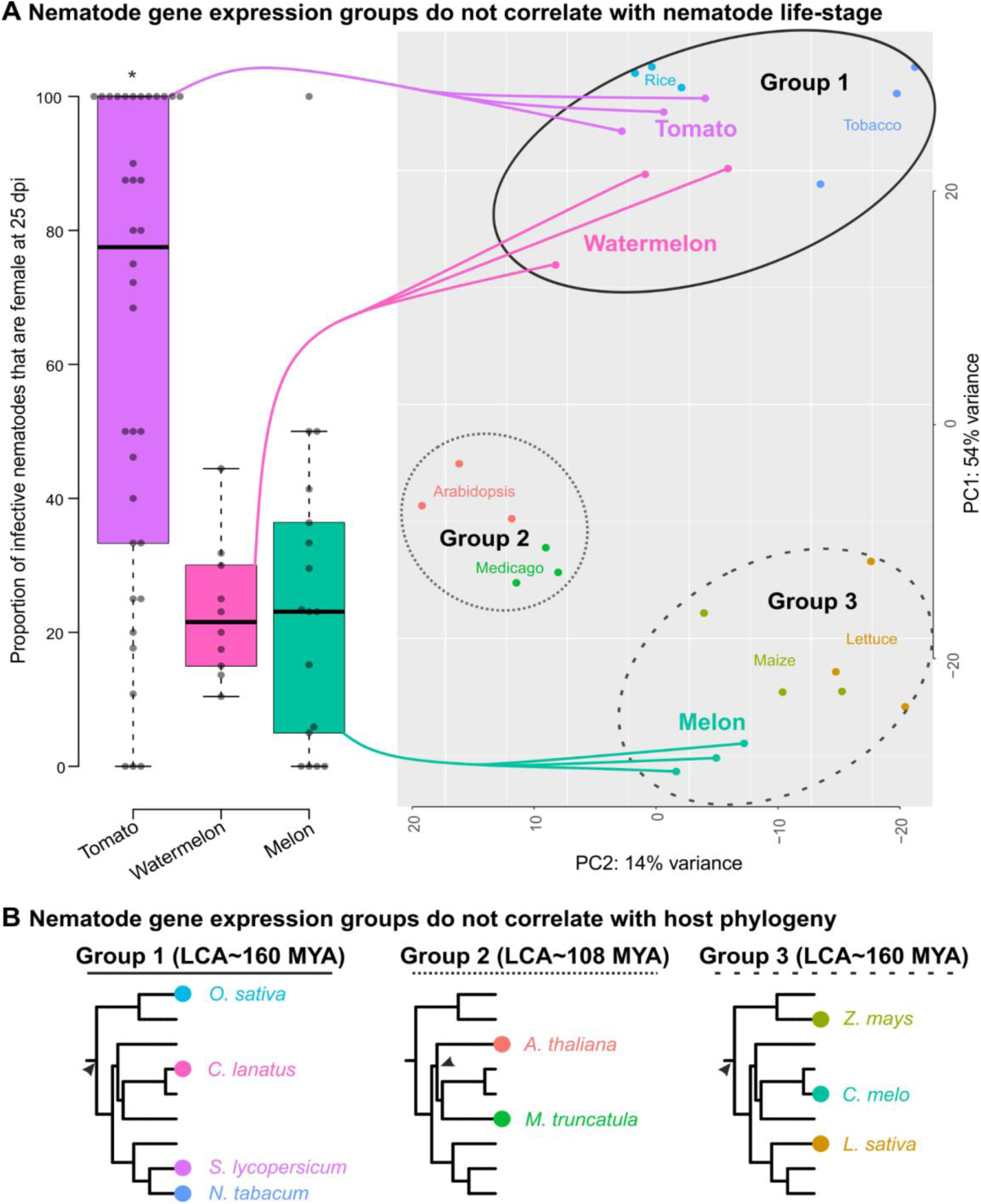
*Meloidogyne incognita* gene expression groups. **A**) Principal components 1 and 2 for *M. incognita* gene expression, showing the three transcriptional programmes. Proportion of female nematodes at 25 DPI for tomato (Group 1), watermelon (Group 1) and melon (Group 3) (Asterisk indicates Dunn’s test, p < 0.001). Centre lines show the medians; box limits indicate the 25th and 75th percentiles as determined by R software; whiskers extend 1.5 times the interquartile range from the 25th and 75th percentiles, outliers are represented by dots. **B**) Host phylogeny (adapted from Phylot v2 (https://phylot.biobyte.de/)) by group. Arrows indicate the Last Common Ancestor (LCA) in estimated Million Years Ago (MYA).

Interestingly, nematode transcriptional groups also cannot be well explained by nematode phenotypes. For example, some of the largest differences in the proportion of nematodes developed to females are from those infecting hosts in the same group (c.f. tomato and watermelon in group 1, Dunn’s test p < 0.05), while some of the smallest differences are from those in different groups (Fig. 2). Taken together, the global transcriptional response of the nematode at 25 dpi are inconsistent with host phylogeny and nematode phenotype (Fig. 2).

### Homeolog triplets contribute to the range of host-specific gene expression

To understand whether and how the hybrid origin of *M. incognita* contributes to host-specific gene expression, we first identified *M. incognita* genes differentially regulated on each host compared to the pre-infection control. This analysis was performed on each host, and a non-redundant list of genes which were differentially regulated during infection of at least one host was generated. To focus on the *in planta* specific predicted secretome, we selected genes that had a mean centred normalised expression value in pre-parasitic J2 < 0.3 and that encoded proteins with a predicted signal peptide for secretion and no predicted transmembrane region (SP and no TM). This resulted in 1,435 putatively *in planta* secreted proteins, ∼6.5% of which have significant sequence similarity to effectors (≥85% identity on BLAST and E-value < 1 x 10^-20^) and were therefore considered the subset which encode putative effectors. To determine the similarities in expression profiles of these genes, a transcriptional network was computed (distance correlation coefficient > 0.96). The majority (67%) of genes in the network are part of one large supercluster (Fig. 3A), with a further 8% having a connection to another effector but not in the supercluster, and the remaining 25% without any connection (Fig. S2).

**Figure 3.**
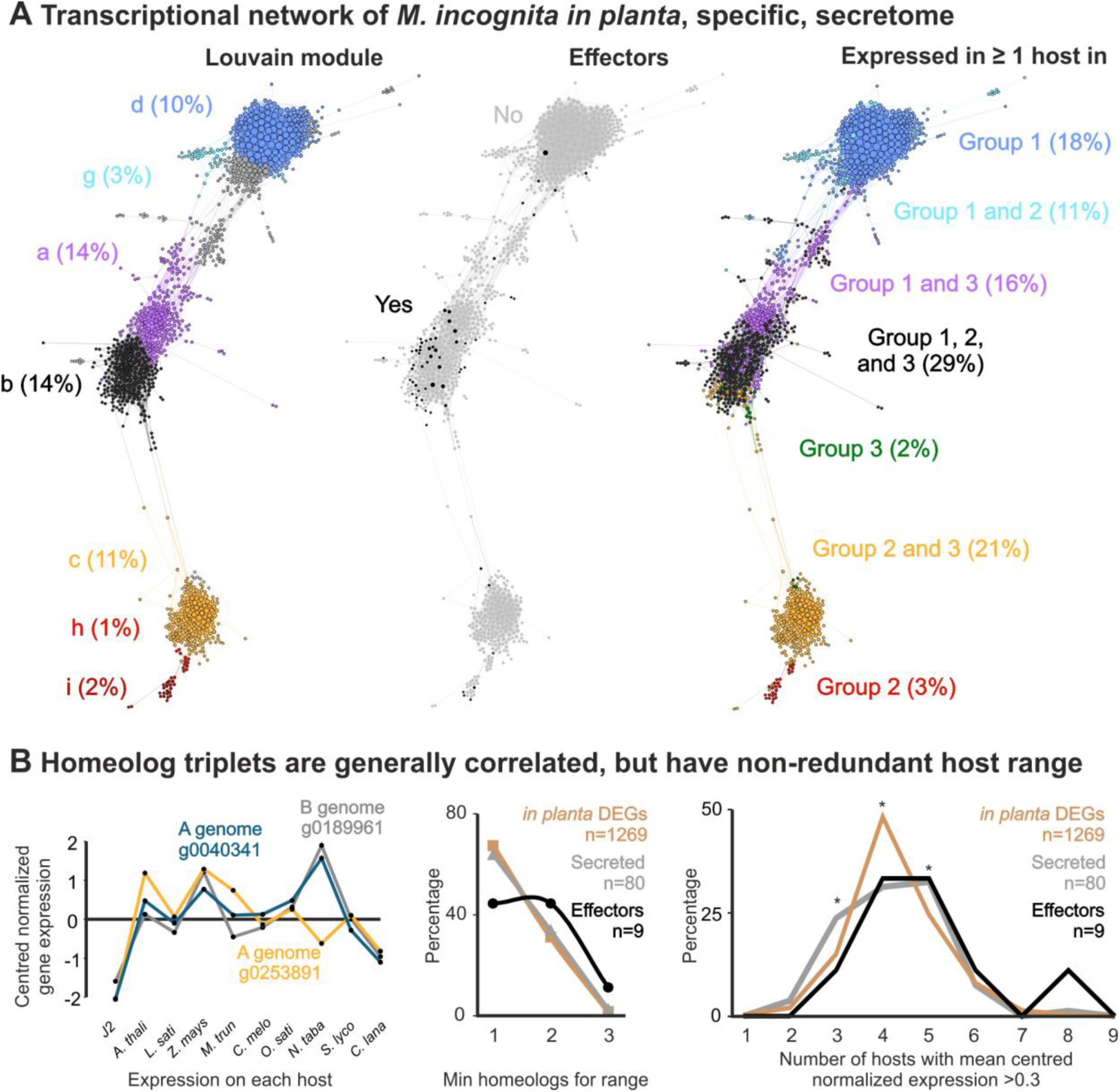
The contribution of homeolog triplets to host-specific gene expression. **A**) Transcriptional network of *M. incognita in planta,* specific, secretome (as defined by DEGs with mean centred normalised expression value in pre-parasitic J2 < 0.3). Each node is an *M. incognita* gene, and lines between nodes indicate a correlation coefficient >0.96. The most connected supercluster (full networks in Fig. S2, and upset plots in Fig. S3) is shown with three colour schemes: left, Louvian modules; middle, putative effectors; and right, statements about expression in groups. **B**) An analysis of homeolog triplets. Left, an example triplet of genes; Middle, the minimum number of homeologs to recapitulate the expression host range of the triplet; Right, number of hosts with mean centred normalised expression >0.3. Asterisk indicates significant differences between *in planta* DEGs and those which are predicted to encode secreted proteins. Secreted protein triplets have a broader distribution of the number of hosts in which they peak expression (more with 3, fewer with 4, and more with 5, Fig. 3B, Hypergeometric test, p = 0.012, 0.0006, 0.025, respectively).

Calculating the Louvain modularity^14^ allows for the non-biased identification of co-expressed clusters within the network (Fig. 3A). The most numerous 9 Louvian modules include the dominant majority of genes in the super cluster. We are able to formalise a definition of this modularity based on the expression of each gene during infection of the three host groups, with high fidelity. For example, Louvian module d (10%) is almost exactly defined as genes with mean-centred normalised expression >0.3 during infection of ≥1 hosts from Group 1 only (Fig. 3A): most often more than one group 1 host (96%), but rarely all four (6.8%, Fig. 3B). Similar comparisons can be made with 7 of the top 9 Louvian modules (Fig. 3A), however, group-exclusive modules are the exception not the rule - only 23% of genes in the network can be described in this way. The majority (77%) are expressed in ≥1 host from ≥1 group. Predicted transcription factors (TF) exhibited a similar pattern. Upon close examination, differentially-expressed TFs were not specific to a single host (except for one in rice), but rather overlapped between multiple hosts. Group 1 exhibited a surprisingly high number of Group 1-exclusive TFs, while Group 2 presents a negligible number, and Group 3 lacks exclusive TFs.

Most putative effectors which are part of the supercluster are in Louvian module b (62%), which broadly aligns with those expressed in at least one host from each group (i.e. mean-centred normalised expression >0.3 on at least one representative from Groups 1, 2, and 3). This is notable because module b contains just 14% of genes, but 62% of putative effectors, in the analysis. The patterns observed for putative secreted proteins indicate that there is a very limited core secretome that exhaustively and exclusively represents a given group. Moreover, these data suggest that expression of the *M. incognita* secretome in general, and the *M. incognita* predicted effectorome in particular, are not deployed uniformly across hosts - but rather on multiple, different, and at present unintelligible combinations of hosts.

To determine whether the homoeologous gene copies, resulting from the hybridisation events (two A subgenomes and one B subgenome), have redundant or different expression profiles across hosts, we identified unambiguous triplets (MCScanX duplication depth = 2 (corresponding to 3 copies), no additional tandem duplicates, n=5167 triplets). Of these, 3,427 triplets contain two or more differentially expressed genes, and of those, 1,269 are *in planta* specific (i.e. mean centred normalised expression value in pre-parasitic J2 < 0.3 in all 3).

Generally, homeologs are concordantly expressed across hosts (e.g. Fig. 3B): 78% of homeologous triplets have a minimum pairwise Pearsons correlation between any two genes above 0.75. To measure the contribution of gene copies to the host range, we calculated the minimum number of homeologs required to recapitulate the expression host range of the sum of the three genes in the triplet. Using this metric, most of these triples (70%) are redundantly expressed across hosts (i.e., the minimum number of homeologs to recapitulate the expression host range of the triplet = 1), while for 29% two homoeologs are required, and for 1% all three are required.

It appears that the two A or the one B subgenomes do not disproportionately contribute to this because: i) computing which gene of the triplet is least similar to the other two copies shows that almost exactly a third are from the B genome (30%), and two thirds from one of the two A genomes (66%); ii) examining triplets where the minimum number of homeologs to recapitulate the host range of the triplet is two (n = 395), also almost exactly a third come from the B genome (31%); and iii) examining genes upregulated on each host, almost exactly a third come from the B genome (min 35.4%, max 37.2%, SD 0.72). Therefore, for approximately one in three cases, homologous triplets have a larger host range expression than the individual genes within, likely as a contribution from all three genomes roughly evenly.

Comparing the subset of these genes which may be involved in the interaction with the plant (i.e. triplets exclusively containing putative secreted proteins (n=80), or those also containing putative effectors (n=9)) reveals a similar picture. However, secreted protein triplets have a broader distribution of the number of hosts in which they peak expression (more with 3, fewer with 4, and more with 5, Fig. 3B, Hypergeometric test, p = 0.012, 0.0006, 0.025, respectively).

### Consequential differential effector expression across susceptible hosts

Given that putative effectors are not deployed uniformly across hosts, we sought to understand the contribution of differential effector expression to host range. Using the same defined proxy for relatively high expression in a given host, compared to all others (mean-centred normalised gene expression above 0.3), most putative effector genes are upregulated in 3 or 4 different hosts (min1, max 6, Fig. 4A). To determine whether differential effector expression can be consequential, Minc18636 was interrogated. This gene was selected because while Nguyen et al. 2018 showed no loss of virulence on tomato following silencing, the transcriptional data generated herein show that Minc18636 is relatively lowly expressed in tomato, when compared to Tobacco, Maize, Medicago, and Rice (in that order, Fig. 2B). To test the differential impact of silencing Minc18636, two hosts were selected: i) Tomato, principally due to the work of Nguyen et al., ^15^ supporting the lack of a role in infection; and ii) Medicago, due to a combination of expression difference (not the highest not the lowest, Fig. 4C), phylogenetic distance (not the furthest not the closest, Fig. 4B), expression group (not the same group as Tomato, but not the furthest (Fig. 2A), developmental progression (broadly similar, Fig. 1), and technical tractability.

**Figure 4.**
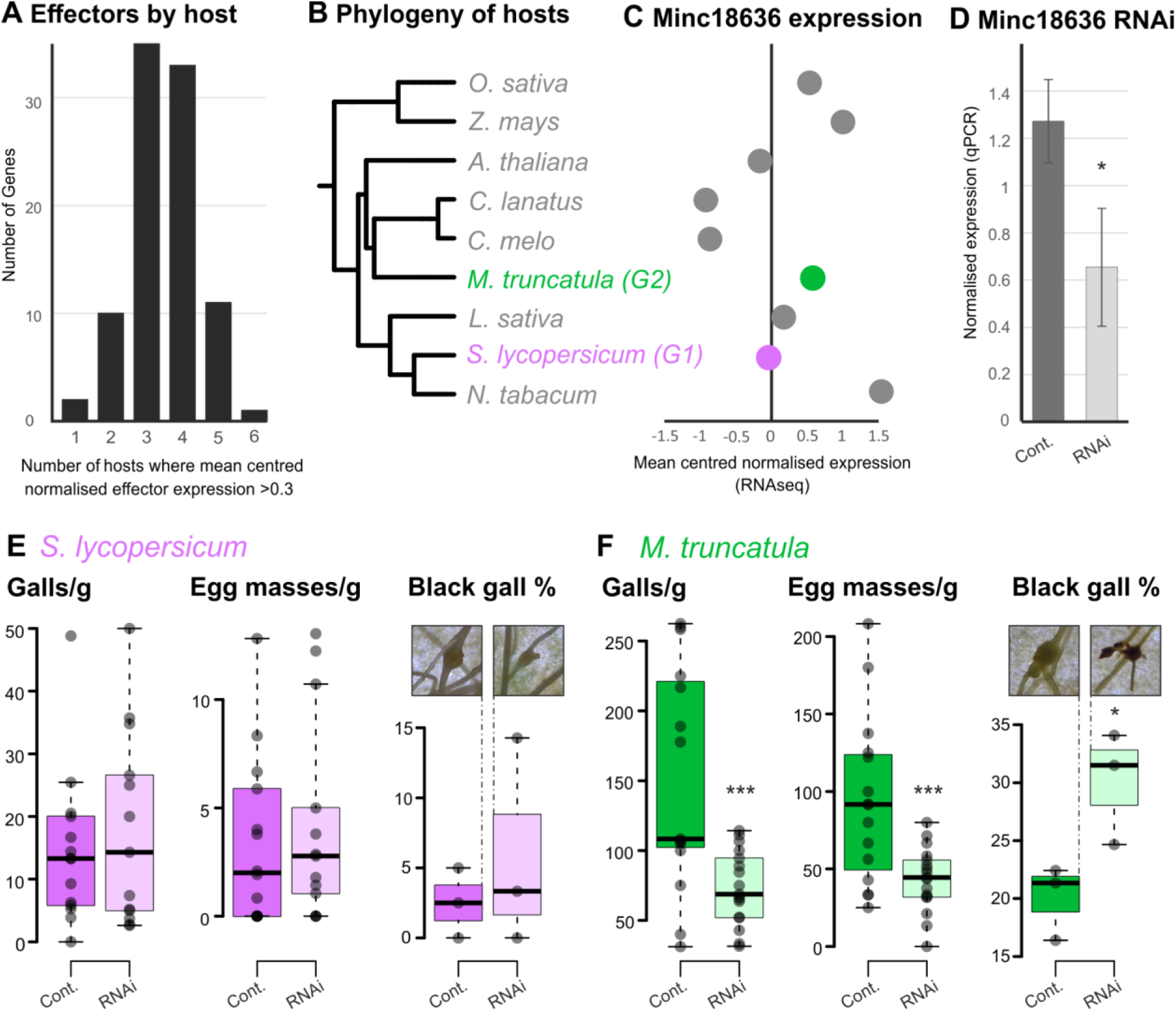
Consequential differential gene expression of *Meloidogyne incognita* putative effector Minc18636. **A**) Frequency distribution of the number of hosts where effector expression “peaks” (mean-centred normalised expression >0.3). **B**) A phylogeny of hosts (adapted from Phylot v2 (https://phylot.biobyte.de/)), with a Group 2 (G2) host and a Group 1 (G1) host highlighted. **C**) Expression of Minc18636 across hosts. **D**) qPCR of Minc18636 following RNA interference (RNAi). Asterisk indicates a significant difference (Student’s T-test, n=3, p-value: < 0.05). Error bars indicate standard deviation. **E** and **F**) Box and whisker plots of the number of Galls per gram of root, Egg masses per gram, and the proportion of galls that were black are shown for *M. incognita* J2s infecting *S. lycopersicum* (left) and *M. truncatula* (right). For black gall assessment, each datapoint represents the mean of five plants originating from one biological replicate of nematodes (n=3), Centre lines show the medians; box limits indicate the 25th and 75th percentiles as determined by R software; whiskers extend 1.5 times the interquartile range from the 25th and 75th percentiles. Significant differences are indicated by asterisks (Student’s T-test, n= 15, p-value: * < 0.05 and ***< 0.001).

Silencing Minc18636 by approximately 48.5 % (Fig. 2C) resulted in no measurable effect on tomato parasitism, largely confirming the previous report, but two notable impacts on Medicago parasitism (Fig. 2D and 2E). Most notably, Medicago inoculated with Minc18636-silenced J2s presented fewer galls per gram of root (−51.2%, Student’s T test p < 0.001) and fewer egg masses per g of root (−53%, Student’s T test p < 0.001). On tomato, however, silencing had no impact on these phenotypes (+23% and +27%, Student’s T Test, p = 1, respectively). It was also noted that Minc18636-silenced J2s infecting Medicago resulted in 50% more dark coloured galls than the control (Student’s T test p < 0.05), whereas Minc18636-silenced and control J2s infecting tomato were indistinguishable (Fig. 4, and Fig. S4).

### There is no “core” gall transcriptome

Finally, to measure similarities and differences in host transcriptional responses to infection, gall-specific gene expression was calculated for each host. These data show that, on average, 12% of the host transcriptome is nematode-dependent at 25 dpi (Table S1, range 2-20%). Typically (in 6/9 hosts) most differentially expressed genes are repressed, with the greatest difference being on Maize (Group 3) and Tobacco (Group 1): in each, approximately twice as many nematode-dependent genes are down-regulated, as up-regulated (6.2% vs 3.2% and 9% vs 4.7% respectively). With the notable exception of tobacco, all other group 1 hosts have more upregulated genes than down, and all hosts with more upregulated than down are in group 1 (from *M. incognita* gene expression point of view). A general case of repression of host gene expression in the galls is consistent with the literature^16^.

To determine the evolutionary history of these differentially expressed genes, we cross-referenced these data with an orthologous gene clustering analysis. Focusing initially on those that arose very recently (i.e. species-specific orthogroups) or expanded very recently (i.e. for each species, where it contributes the most genes to an orthogroup and contributes more than three times the next highest contributing species), a similar trend of more repression than activation is observed (Fig. 5). To compare genes with an opposite evolutionary history, i.e. those which likely have an ancient origin and are conserved to the present day, we identified orthogroups with at least one representative from each species (8,524 OGs). While just 32% of these orthogroups contain no genes differentially expressed in any host, of the remaining 5,729 orthogroups, the dominant majority (71%) are consistently differentially expressed - meaning, all differentially expressed genes, from any species in the orthogroup, are up, or down, but not a mix of both. The first striking observation is that in these common and consistently differentially expressed genes, we find an opposing trend to genes as a whole: 58% contain upregulated genes, while 42% contain down-regulated genes. These orthogroups contain almost the same number of genes (4,779 and 4,815, respectively) but includes a greater diversity of functional categories (317 up vs 231 down). While we have a general pattern of down-regulation of all genes, we have a general pattern of up-regulation of the greatest diversity of conserved genes.

**Figure 5.**
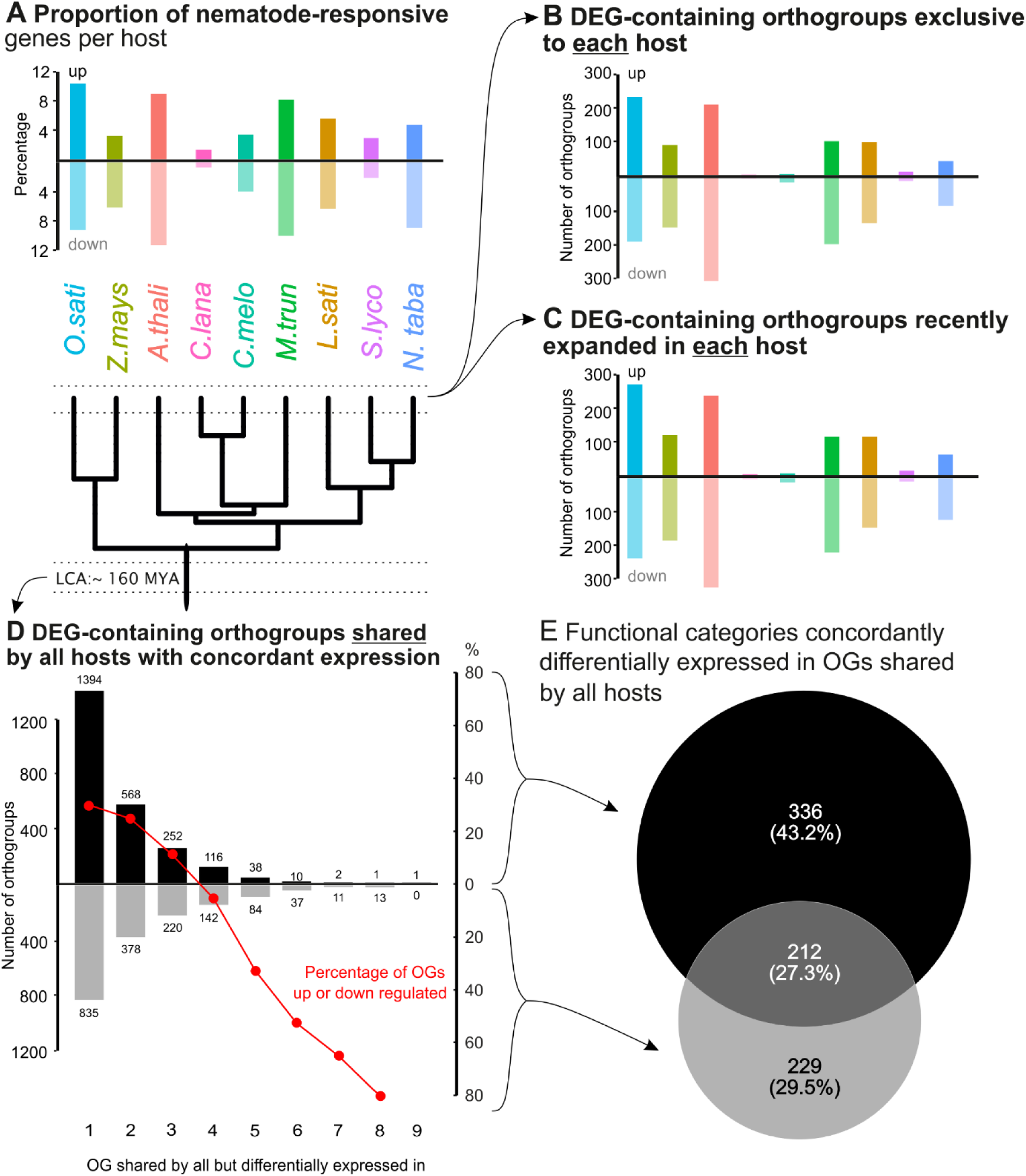
Contrasting expression patterns of apomorphic and conserved nematode-responsive plant genes. **A**) Percentage of nematode-responsive differentially expressed genes (DEGs) per host species. **B**) Number of species-specific Orthogroups (OGs) with > 0 nematode-responsive DEGs concordantly expressed (i.e. all DEGs in OGs are up, or down, but not a mix). **C**) Number of species-expanded Orthogroups (OGs) with > 0 nematode-responsive DEGs concordantly expressed (i.e. all DEGs in OGs are up, or down, but not a mix). **D**) Number of orthogroups shared by all hosts, which have > 0 nematode-responsive DEGs unambiguously up, or down, regulated in one to nine hosts. Red line indicates the percentage of these OGs which are up or down (right axis). **E**) Overlap of functional categories (Gene Ontology terms) in OGs of panel D (shared by all hosts, which are unambiguously up- or down-regulated in one to nine hosts).

Interestingly, this difference is largely driven by genes which are common to all species, but only nematode-responsive in a small subset of hosts. Those orthogroups shared by all hosts but only consistently differentially regulated in 1, 2, or 3 hosts are dominated by up-regulation, while those orthogroups shared by all hosts but consistently differentially regulated in 5, 6, 7, or 8 hosts are dominated by down-regulation. Essentially, most individual species (7/9) follow this trend (Fig. 5). Given that it is more common to find orthogroups shared by all hosts but only differentially regulated in a few of them, this subset skews the pattern of the whole. These data also highlight a striking observation: there is only a single orthogroup shared by, and concordantly up-regulated in, all species (OG0001156: Heavy metal-associated isoprenylated plant proteins), which was recently associated with susceptibility in *A. thaliana* to *M. incognita*^17^. There are zero orthogroups shared by, and concordantly down-regulated, in all species. Taken together, we conclude there is no “core” gall transcriptome at 25 days post-infection.

## Discussion

While we have known for some time that *M. incognita* changes gene expression across its life cycle^18^, these data show that at a given point in time (25 days post-infection), *M. incognita* has distinct transcriptional signatures depending on the host it infects. Broadly speaking, three transcriptional groups were identified across the 9 hosts tested. The lack of universal congruence between host phylogeny, host physiology, and nematode developmental stage with nematode gene expression groups is puzzling. Regarding nematode stages, 3 out of 4 members of group 1 present the highest percentage of mature females, whereas other hosts present similar proportions but are in different groups (e.g., watermelon-melon and maize-rice). We cannot rule out that this pattern is determined by an unmeasured phenotype (e.g. polysaccharide composition of the roots), or that it results from other poorly-characterised factors, such as the history of host domestication, or the historically predominant hosts of *M. incognita* at its geographical origin. Such historical factors might affect the host-parasite coevolution, and ultimately, the complexity of the transcriptional signature for each host. Taking Group 1 again as an example, tomato and tobacco originated in South America, as well as many wild rice (such as *Oryza latifolia*). Taking all these observations together, it seems likely that several factors contribute to this complex system, but that none is dominant.

Despite the fact that the underlying reasons for the observed differences in nematode gene expression are at present unintelligible, these differences in nematode gene expression clearly can matter. We demonstrate this by interrogating the role of effector Minc18636, which is differentially expressed on some hosts but not others at 25 dpi. We show that J2s silenced for this gene form fewer and aberrant galls on a host on which it is highly expressed, when compared to a host on which it is lowly expressed. Clearly, then, the upregulation of certain effectors on certain hosts can be consequential. While this is the first demonstration of consequential host-specific effector expression for plant-parasitic nematodes, caution should be taken with extrapolation. It seems plausible that other genes differentially expressed on some hosts but not others are similarly required. However, until more examples are described, causal relationships should not be assumed. The discovery of a causal relationship herein was made possible because the work of Nguyen et al.^15^, which reported the absence of a phenotype when silencing this effector on tomato. This highlights the importance of reporting negative results for the progress of the field at large.

A long-standing idea in the field is that the polyploidy of *M. incognita* may contribute to the remarkably wide host range. Preliminary analysis showed that homeologous gene copies emerging from allopolyploidisation frequently display different gene expression patterns during the developmental life cycle of *M. incognita*^11^. However, no direct connection to the host range could be done at this time. The first challenge is to define which multi-copy genes likely arose from the ancient hybridisation events (as opposed to subsequent duplication events). *M. incognita* has a triploid genome made of two relatively close AA’ sub-genomes and a more distant B sub-genome^19,20^. Therefore, we focused on collinear syntenic genomic regions that remained in exactly three copies as a proxy. We excluded all the other cases that might include extra duplications or copy losses, as they might result from other independent duplication events. While there are other explanations for the presence of such triplets (e.g. one from each of two hybrid parents but nothing from the third with a subsequent duplication of one copy), the most parsimonious explanation is that they arose from the hybridisation events and have been maintained to the present day. In each triplet, we can determine which came from the two A sub-genomes, and which came from the B sub-genome, based on the contig it resides in^20^, but cannot distinguish between the As due to their equal symmetric distance to B. With the caveats described, we think this is a good proxy for genes of hybrid origin. The majority of these gene copies have highly similar expression profiles across the hosts tested here. However, for approximately 30% of the triplets (thousands of genes), the expression host range of the triplet is larger than the expression host range of any individual gene within the triplet.

Taken together, we therefore conclude with confidence that the hybridisation events have contributed to a greater expression host range for this species. What is less clear is the pathway to this contribution. Curiously, despite the higher nucleotide sequence divergence of the B subgenome compared to the A subgenomes; the B subgenome does not contribute more to the expression divergence between gene copies within the triplets. Yet, this observation is consistent with the absence of a systematic genome dominance observed for either the A or the B subgenomes^20^. So far, we do not know, nor can we determine, the ancestral state of each gene from the parents of the hybridisation events, because they are either extinct or not yet identified. Therefore, it may be that the expression pattern of these genes in the hybrid parents was different from one another, and that the extant triplets in *M. incognita* simply reflect the sum of the parents. However, it may also be that the expression pattern of these genes in the hybrid parents was not different from one another, and that following hybridisation subsequent diversification/neofunctionalization has occurred. For the latter, one homeolog may have lost expression on a given host, or one may have gained expression on a given host - both would manifest as a greater summed host range of the homologous triplet than the individual genes within. In the absence of evidence, it seems plausible that both scenarios contributed to the present-day observation that homologous triplets have a greater range of expression on hosts than the individual genes themselves in approximately 30% of cases.

Investigating the plant side of interaction, we find that even though the total number of nematode-responsive genes at 25 dpi can vary across an order of magnitude (2-20% of the plant genome), they tend to respond in broadly similar ways during infection. For example, for most hosts, this substantive differential expression is skewed towards suppression, which is consistent with the literature and the proposed link to methylation^16^. Interestingly, genes with opposing evolutionary histories also have opposing expression trends. Genes that evolved or expanded recently are primarily down-regulated, while genes that have an ancient origin and are conserved to the present day are primarily up-regulated. Together, these paint a picture of general gene repression, but widespread activation of core conserved genes. Activation of highly conserved genes has some parallels to the patterns observed for *H. schachtii* infection of *A. thaliana* - lineage-specific nematode genes (including, effectors) modulating the expression of widely conserved plant genes^21^. However, these analyses paint only a partial picture because while it is true that most *M. incognita*-responsive host genes in orthogroups conserved in all species are concordantly expressed, not all genes in those orthogroups are differentially expressed. In fact, the majority contain differentially expressed genes on only a small number of hosts, with the remaining genes in the orthogroup being non-nematode responsive. Put in another way, in spite of abundant conserved orthogroups, there is only a single concordantly upregulated orthogroup in all hosts at 25 days post-infection. This finding means that extrapolating from one host, even to close relatives, will almost certainly be in error - posing a major challenge to the generalisability of susceptibility genes discovered by transcriptomic analyses^21^.

At the time of sampling, 25 days post-infection, nematodes have successfully established long-term biotrophic interactions with the host. Therefore, presumably the galls, and certainly the giant cells within, must fulfil the same/similar function with respect to the nematode. They must be suppressed in their immune response and provide sufficient nutrition to the nematode development and egg production. Yet, the fact that they are so transcriptionally distinct leads us to consider two, not necessarily mutually-exclusive, models: i) There are many possible routes to a functioning gall (i.e. “all roads lead to Rome”); or ii) late-stage gall transcriptomes are dominated by effect, not cause.

## Methods

### Obtaining plants

Plant hosts were selected based on a series of criteria to aid understanding and discovery. Firstly, they must be susceptible to *M. incognita*. Secondly, they must be able to grow under the same conditions. Thirdly, genomic/transcriptomic resources must be available. Lastly, and equally important, we sought to have representatives not only across the phylogenetic tree of flowering plants, but also some hosts from the same family. Therefore, nine susceptible hosts distributed in six plant orders were selected (Table S4):

Except for *A. thaliana*, tomato, and tobacco, seeds of all hosts were surface sterilised (70% alcohol and 20% bleach, each for 1 minute, followed by four washes using sterile water), and pre-germinated axenically for 48 h (dark at 30°C). After germination, they were transferred to seed germination pouches (CYG™) and placed vertically in a box containing Hoagland’s nutrient solution in a growth chamber (6h/8h light/dark at 30°C/21°C). For tomato, fresh cuttings were rooted in Hoagland’s solution and subsequently transferred to seed pouches. Regarding *A. thaliana* and tobacco, seeds were grown in Knop medium (21°C for ∼ 14 days), after which they were transferred into the aforementioned growth chamber to acclimate before inoculation. While Arabidopsis were kept on media, tobacco plants were transfer into pouches.

### Nematode acquisition and subculture

*Meloidogyne incognita* ‘Morelos’ culture was started from previously galled roots. To ensure the stability of the population, new inocula were produced by transferring galled roots from tomato ‘Moneymaker’ into 5-week-old plants. The inoculum used in infection experiments consisted of 200 second-stage juveniles per plant. To obtain them, galled roots were removed from pots, cleaned and blended for 1 minute with a 10% bleach solution to release the eggs. The eggs were hatched, and J2 density was estimated by using Peters’ counting slide. Nematodes were pelleted and resuspended using a 0.01% Tween^®^ solution (“Nemawash”). Inoculum volume was adjusted to 1 J2 per μL.

### Inoculation

Plants growing in pouches were laid flat on trays and cut open laterally to expose the roots. Inoculum droplets were then applied to the upper half of the root systems. The pouches were stacked and incubated horizontally in a box containing wet paper. After 48 hours, they were transferred and positioned vertically in a new box with Hoagland’s solution. For *A. thaliana* growing on petri dishes, nematodes were initially cleaned under a dissection microscope by removing debris and adding clean Nemawash. Afterwards, nematodes were transferred to a 50 mL tube and surface-sterilised by soaking in an antibiotic solution (CTAB 0.5 mg/mL (Sigma) + 0.1% v/v chlorexidine digluconate (Sigma), kanamycin 0.1% w/v, 0.6% streptomycin EDTA solution (Sigma) and Tween^®^ 0.01% v/v) for 25 minutes, followed by six washes with sterile water. The cleaned nematodes were then pipetted on *A. thaliana* roots under axenic conditions, incubated horizontally for 48 hours, shifted to a vertical position and maintained in a growth chamber as described previously.

### Sample collection, RNA extraction and transcriptomic analysis

Gall collection was performed 25 dpi. By using a dissection microscope, tweezers and a scalpel, galls were excised from the roots and stored in 2 ml Eppendorf tubes placed on ice. As a control, symptomless and comparable adjacent fragments of roots were collected. Each replicate consisted of a mixture of galls, with the corresponding control fragments added to a matching, numbered control tube. After sampling, the tubes were flash-frozen with liquid nitrogen and stored at −80°C. The total RNA was extracted by adding two metal beads and grinding using Geno/Grinder 2010 (six cycles at 300 rpm and 30 seconds). After the first two cycles, 450 µL of lysis buffer (RNeasy^®^ Plant Mini Kit) + 1% β-mercaptoethanol were added to each tube and subjected to five more cycles. After becoming a fine powder, total RNA was extracted using RNeasy^®^ Plant Mini Kit following the manufacturer’s instructions with a DNase I step. Quality control and quantity were assessed by using Nanodrop™ and by electrophoresing the RNA on a 1% agarose gel at 100 V for 20 minutes.

Pre-parasitic J2s were included as a comparative group for identifying *M*. *incognita in planta* DEGs. For RNA extraction, freshly hatched nematodes were collected and cleaned, as previously described. During this process, other nematodes (i.e., free-living nematodes), *M. incognita* males and sedentary stages were removed. Total RNA was extracted from four samples, each containing 15,000 J2s, using the aforementioned method.

RNA sequence and mRNA library construction were carried out by Novogene (Cambridge, UK). The library was prepared using poly-A enrichment (poly-T oligo-attached magnetic beads), followed by fragmentation, cDNA synthesis (with random hexamer primers), end-repair, A-tailing, adapter ligation, size selection, amplification, and purification. The Illumina sequencing was performed using 150 bp paired-end reads. Sequence volume generated for maize, Medicago and tobacco was 24Gb and 12Gb for galls and control fragments, respectively. A volume of 15Gb was used for all other hosts’ galls, and an 8Gb volume was used for their control roots and J2s samples.

Raw data were analysed using FastQC v.0.11.9^22^. Adaptors were removed and 10 bp were trimmed using BBDuk (BBTools v.38.18 - https://github.com/BioInfoTools/BBMap/blob/master/sh/bbduk.sh). To obtain *M. incognita* DEGs, we used a combination of multi-mapping and RSEM/EBseq based on Kozlowski et al.^23^. Using STAR v. 2.7.10a^24^, trimmed reads were first mapped into the respective plant genome, and the unmapped reads were extracted and mapped to the v.4 *M. incognita* genome^20^. Mapping rates of first and second steps are shown on Table S5.

To obtain host data, the reads of diploid plants (2n) were mapped into a joint plant + *M. incognita* genome, whereas polyploid plant reads (i.e., tobacco) were mapped similarly to *M. incognita* - firstly mapped to *M. incognita* genome and the unmapped reads were extracted and mapped into the tobacco genome. Counts were estimated using RSEM/EBseq v.1.3.1^25,26^, and differentially expressed genes (DEGs) were obtained by comparing galls to pre-parasitic J2s (*M. incognita* DEGs) or comparing galls to control fragments (hosts DEGs) (FDR at 0.05). The EBseq uses the expected counts generated by RSEM and Bayesian modelling to control sample-to-sample variability^26^.

Data was further processed on R 4.2.1.^27^, where Principal Component Analysis (PCA) was carried out using the R Package DESeq2^28^. Afterwards, Log2FC was added to select up- and down-regulated (Log fold change ≥ 0.5 and ≤ - 0.5, respectively). After obtaining up- and down-regulated *M. incognita* genes, hierarchical clustering was performed after scaling using hclust from the stats package v.4.2.1. on R^29^. A schematic overview of this method is described in Fig. S5.

### Nematode development assessments

Nematode development assessments were conducted in two batches: First: lettuce, maize, medicago, melon, tomato, and watermelon; Second: *A. thaliana*, rice, and tobacco. Plants were obtained and inoculated as previously described, using 200 J2s (1 J2/ μL). Twenty-five days post infection, pictures of galls were recorded and nematodes within roots were stained using acid fuchsin^30^ and stored in acidified glycerin (1 mL HCl per L of glycerol). The number of galls, J2s, J3s, J4s and adults (male and female) were counted under a dissecting microscope. Then, nematode/galls were estimated, and the gall ratio (Gall ratio = ⌀Gall/mean (⌀ adjacent roots)) was assessed using the pictures taken. Gall ratio data was transformed using Log (X+1) and normality was checked using the Shapiro-Wilk test (p > 0.05). Then, it was analysed using ANOVA post-hoc Tukey’s test using the Laercio R package v. 1.0-1^31^. For non-normally distributed data, they were analysed using dunn.test R package v.1.3.6^32^ through the Kruskal-Wallis test post-hoc Dunn’s test.

### Transcriptional network analysis

To assemble the transcriptional network, we initially predicted *Meloidogyne incognita* secretome by selecting proteins presenting a signal peptide and not bearing a transmembrane domain, which were obtained using SignalP 4.1^33,34^ and TMBed 1.0.0^35^. To define expression on a given host, mean centred normalised gene expression > 0.3 in a host was considered a “Peak”; if 0, “Neutral”; and if < 0 “Trough”. *M. incognita* genes were considered differentially expressed *in planta* if value in pre-parasitic J2 < 0.3, further classified as “Group 1”, “Group 2” “Group 3”, or combinations of, if > 0.3 in at least one representative of the PCA host groups (Group 1 = rice, tobacco, tomato and watermelon; Group 2 = Arabidopsis and Medicago; Group 3 = lettuce, maize and melon). A curated list of putative and known effectors available in the literature for *M. incognita* was used to identify effectors in our network (Kika et al., unpublished and ^36^). Their amino acid sequences were compared to the *M. incognita* v.4 predicted proteome using BlastP 2.11.0+^37^ and it was considered a match if identity was higher than 85% and E-value < 10⁻²⁰. The network was generated following Molloy et al. instructions and custom scripts^38^, in which data was loaded into R using the energy R package^39^, and pairwise distance correlation between genes was calculated at an arbitrary edge threshold of 0.96. Visualisation and Louvian module calculations were performed using Gephi 0.10.1^40^.

### Transcriptional factor analysis

AnimalTFDB4^41^ was used to identify putative transcription factors from the predicted protein sequences of all *M. incognita* genes. In subsequent analyses, we excluded transcription factors that were not differentially expressed (< 0.3), or that were differentially expressed in pre-parasitic J2s as compared to controls, as is consistent with the other analyses above.

### Orthogroup analysis

To assemble host orthogroups, predicted proteomes from each respective plant were processed using OrthoFinder 2.5.5 using default parameters. A custom script was used to process the output further to highlight the number and identity of corresponding differentially expressed genes. Gene ontology (GO) terms were added to each orthogroup using InterProScan version 5.66-98, with specific parameter settings as ‘-f tsv --appl Pfam -- goterms -pa –iprlookup’^37,42^. A total of 36,536 orthogroups were obtained, and the ones shared by all hosts were filtered and classified as “unambiguous up” or “unambiguous down” if the DEGs were unambiguously up- or down-regulated in one to nine hosts. Conversely, we also filtered orthogroups only present in a particular host, or those that were expanded more recently in a given host. Then, groups presenting unambiguously up- or down-regulated genes were selected. Other classifications and custom codes can be found at https://github.com/vmourade/HostRangeParadox and https://github.com/chongjing/OrthoFinder_GO.

### RNAi

For RNAi experiments, Medicago ‘A17 Jester’ and tomato ‘Heinz 1706’ were used, and plants were obtained as previously described. The siRNA primers were obtained from Nguyen et al.^15^, and Minc18636 siRNA was synthesised using Ambion Silencer siRNA Construction Kit (AM1620, Ambion) following the manufacturer’s instructions. The J2s were equally distributed in six tubes and soaked in a siRNA solution containing octopamine (1M) at 0.5%^43^. Control nematodes were treated similarly, but using siRNA targeting GFP. After 24h, nematodes were washed twice using ultrapure water, and plants were inoculated with 120 J2s. Remaining nematodes of each tube were flash-frozen and retained for qPCR confirmation of knock down. Plants were evaluated six week following inoculation, where the number of galls per gram, egg masses per gram, and the proportion of darkened/black galls were counted. qPCR was carried out using three replicates for each treatment, using *GAPDH* and *HK14* as the reference genes for normalisation^15^. Statistical significance of both qPCR results and phenotype evaluations was analysed using Student’s T-test on the R program.

## Acknowledgements

VHMS would like to thank the European Union’s Horizon 2020 Research and Innovation Programme under Marie Skłodowska-Curie grant agreement 101025218. Work on plant-parasitic nematodes at the University of Cambridge is supported by DEFRA licence 125034/359149/3, and funded by BBSRC grants BB/R011311/1, BB/S006397/1, BB/X006352/1, and BB/Y513246/1, a Leverhulme grant RPG-2023-001, and a UKRI FrontierResearch Grant EP/X024008/1, Royal Society research grant RGS\R1\231239, and through the Cambridge-Africa ALBORADA Research Fund. The authors would like to thank Dr Bruno Favery’s research group for kindly providing detailed protocols and primers previously used in Nguyen et al. (2018). Authors also acknowledge the Plant Sciences Teaching laboratory team, Dr Doris Albinsky, Dr Jordi Garcia, Dr Katharina Schiessl, and Prof. Uta Paszkowski’s research group for providing seeds used in our work. The authors would like to thank Mr. Junior Lusu Kika for providing the list of effectors, as well as Roberta Healey and Dio Shin for their assistance in our work. We would also like to thank Dr Marc Bailly-Bechet for his guidance on PCA comparisons.

---

**Figure S1.**
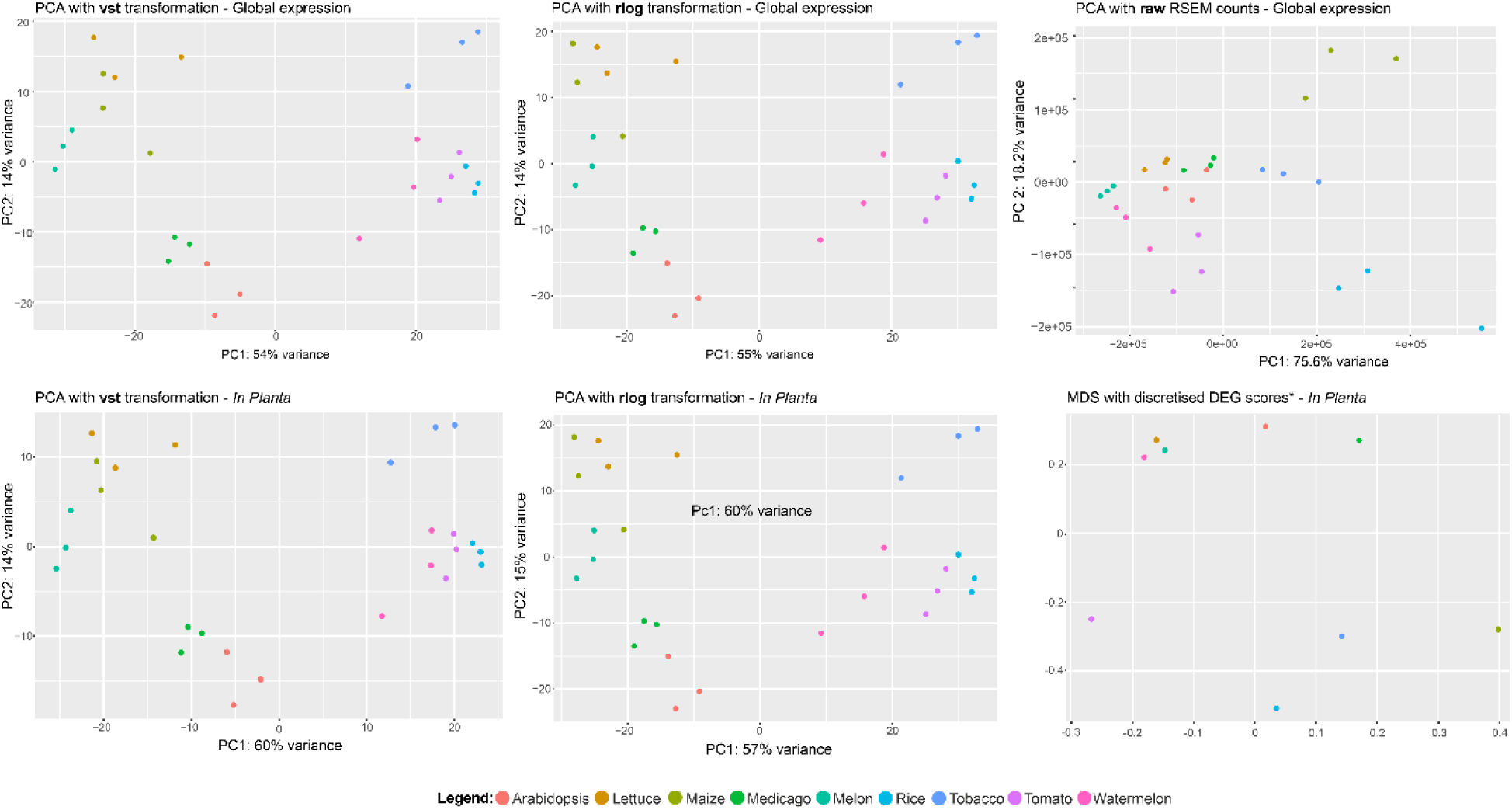
Inconsistent grouping of nematode gene expression by host. Various dimensional reductions (PCA or MDS) are shown, for nematode gene expression on each of 9 hosts (coloured the same for each plot).

**Figure S2.**
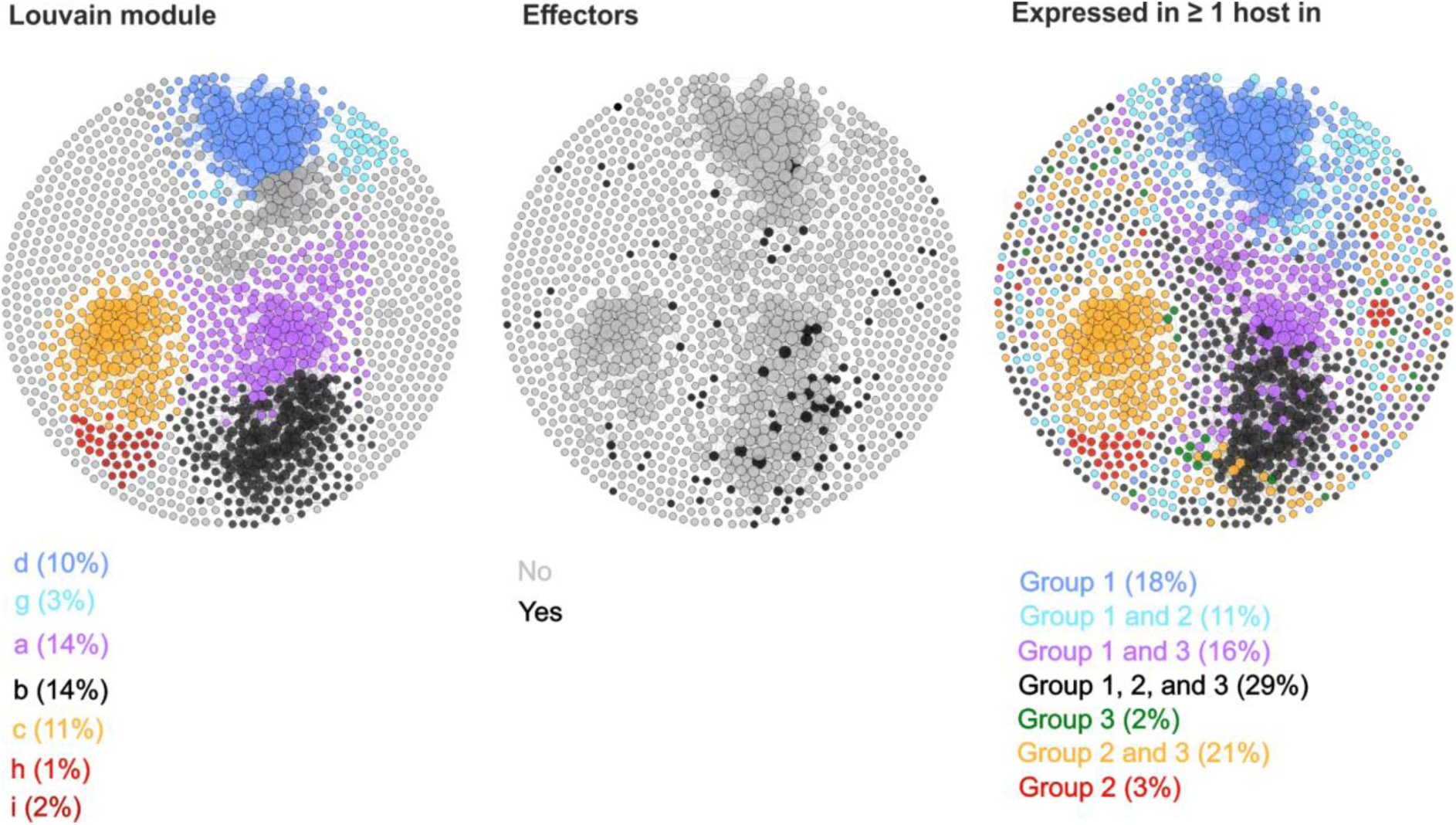
Transcriptional network of *Meloidogyne incognita in planta*, specific, DEGs. Transcriptional network of *M. incognita in planta,* specific, secretome (as defined by DEGs with mean centred normalised expression value in pre-parasitic J2 < 0.3). Each node is an *M. incognita* gene, and lines between nodes indicate a correlation coefficient > 0.96. The diameter of the node is proportional to the number of connections, and are coloured by: left, Louvain modules; middle, putative effectors; and right, statements about expression in groups.

**Figure S3.**
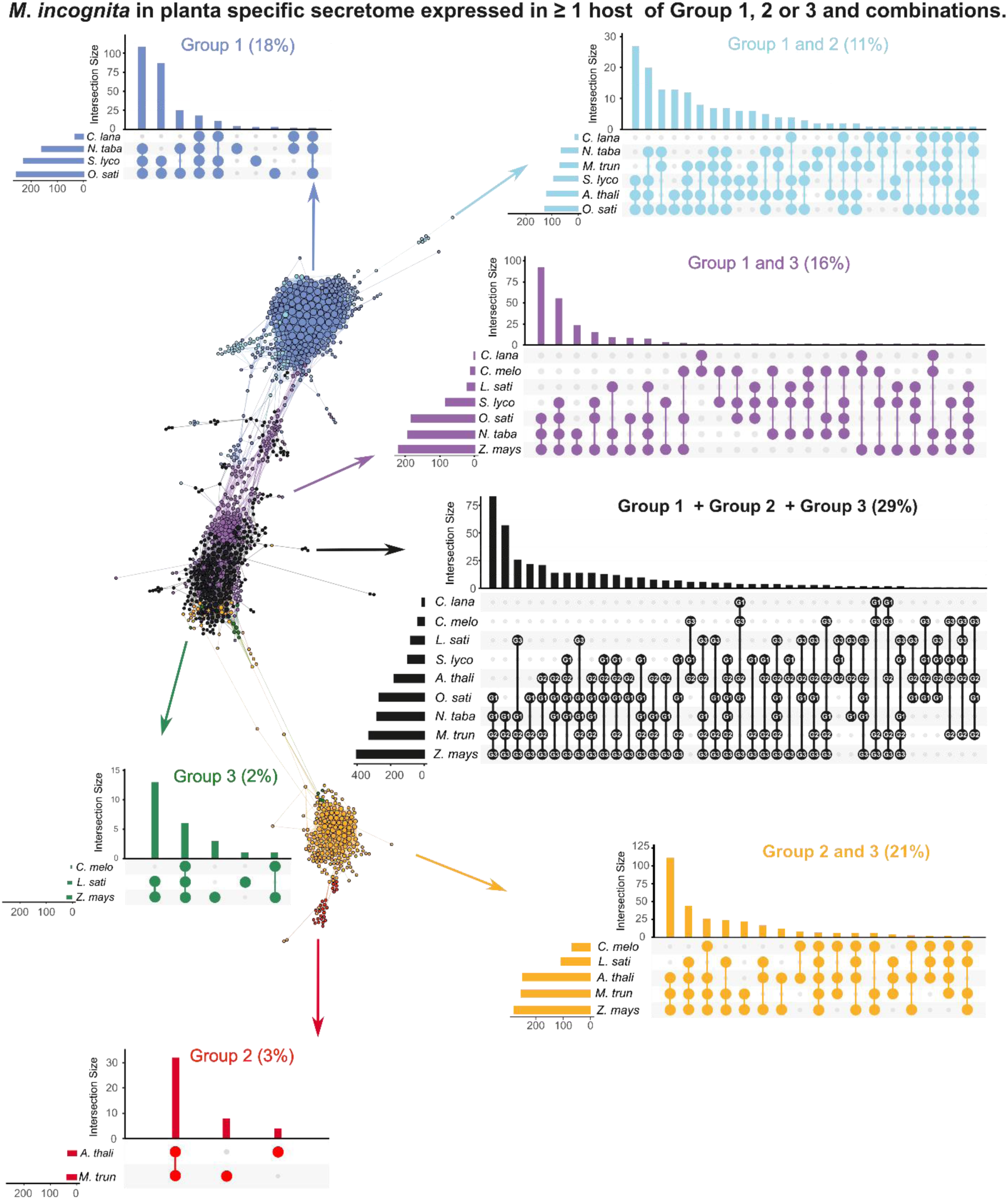
Upset plots for transcriptional network of *Meloidogyne incognita in planta*, specific, DEGs. Transcriptional network of *M. incognita in planta,* specific, secretome (as defined by DEGs with mean centred normalised expression value in pre-parasitic J2 < 0.3). Each node is an *M. incognita* gene, and lines between nodes indicate a correlation coefficient > 0.96. The diameter of the node is proportional to the number of connections, and are coloured by statements about expression in groups. For each category, an upset plot shows the distribution of host gene expression.

**Figure S4.**
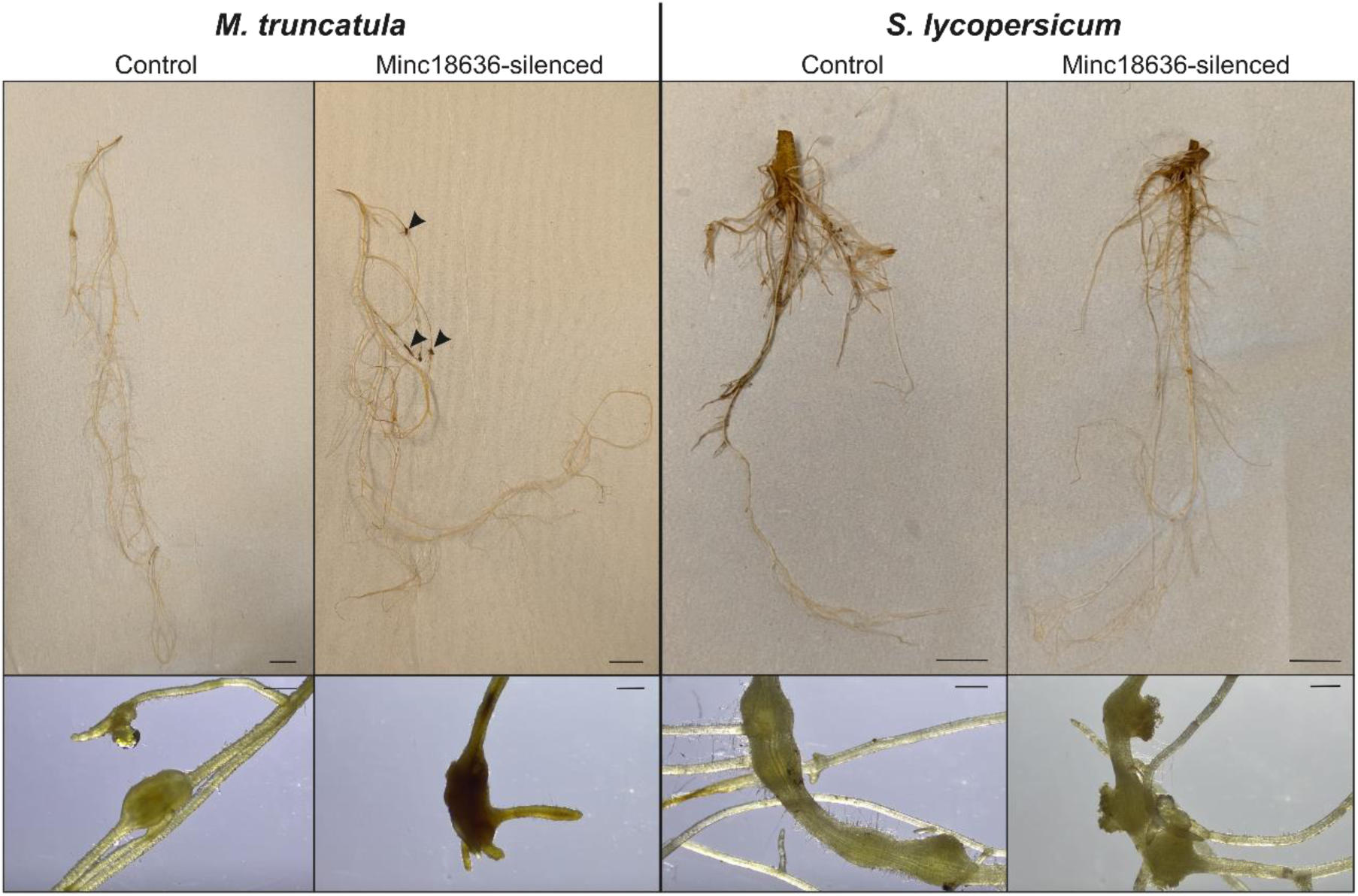
Host phenotype following infection of *Meloidogyne incognita*. Phenotypes of *Medicago truncatula* (left) and *Solanum lycopersicum* (right) roots infected with *M. incognita.* Untreated and Minc18636-silenced J2s are indicated. Arrows indicate dark/black galls. Scale bars of the main image indicate 1 cm, scale bars of the inset indicate 0.1 cm.

**Figure S5.**
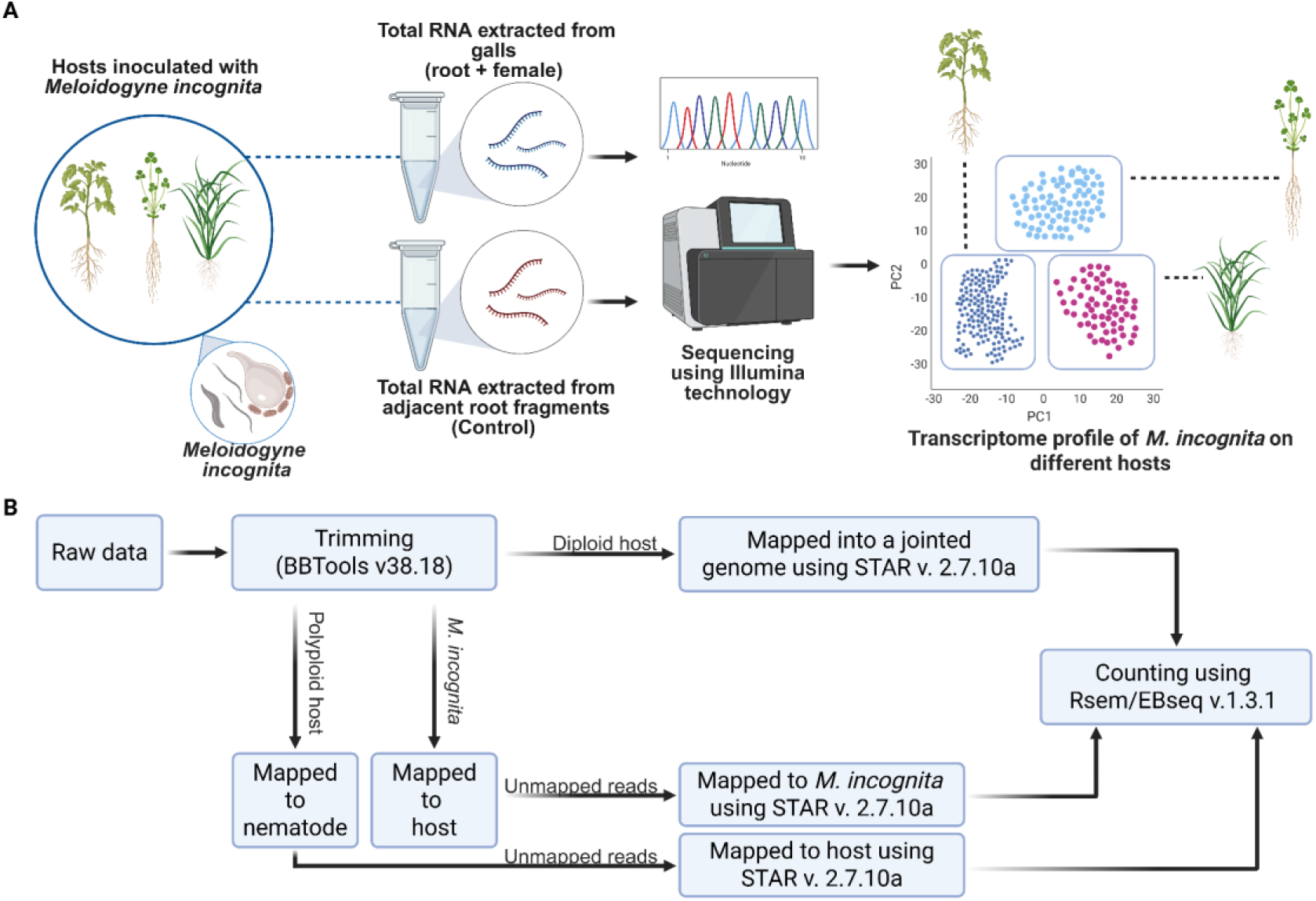
Schematic overview of the present work. **A)** Summary of the experiments. **B)** Bioinformatic pipeline used to process *Meloidogyne incognita*, diploid and polyploid hosts reads. Figure prepared using BioRender (https://www.biorender.com/).

**Table S1.**
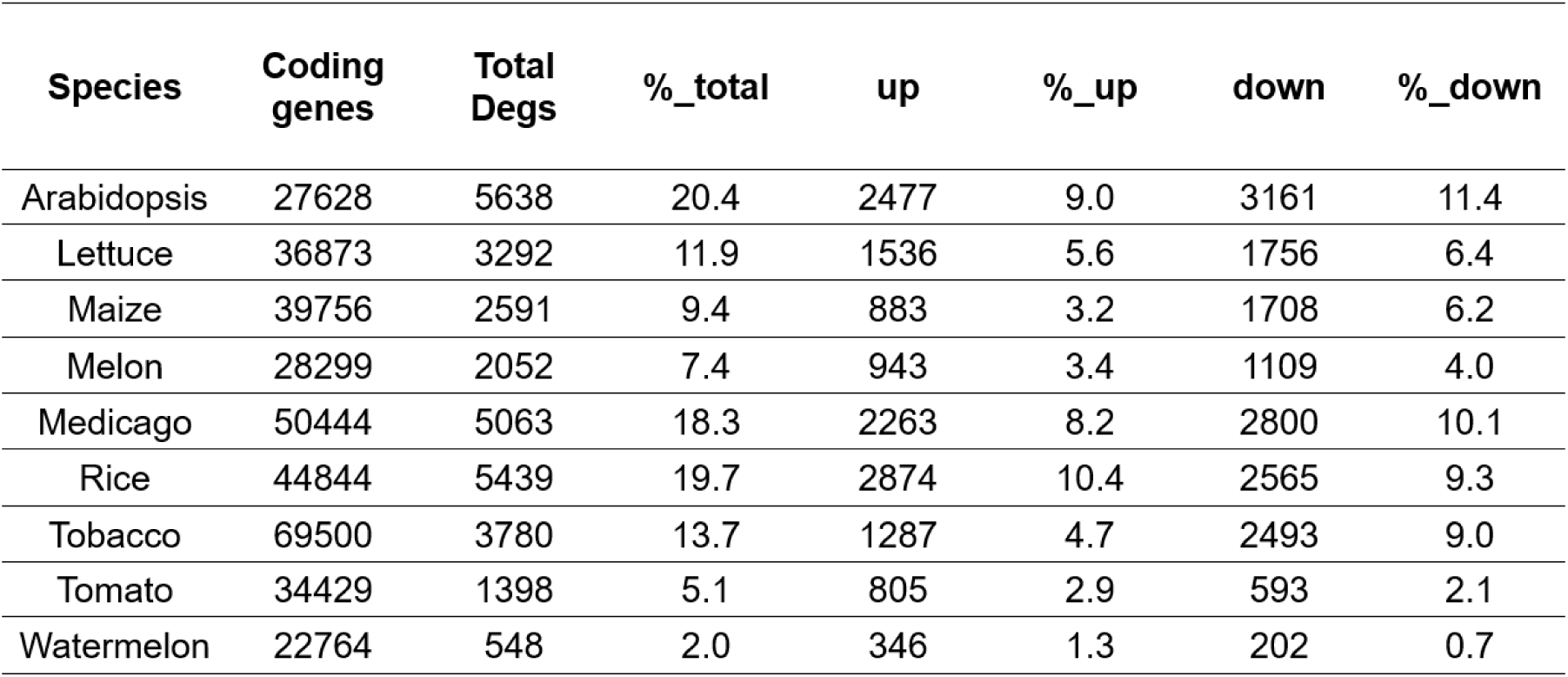
Total number of coding genes for each species, the number of differentially expressed genes (DEGs), counts of upregulated and downregulated genes, and their respective proportions.

**Table S2.**
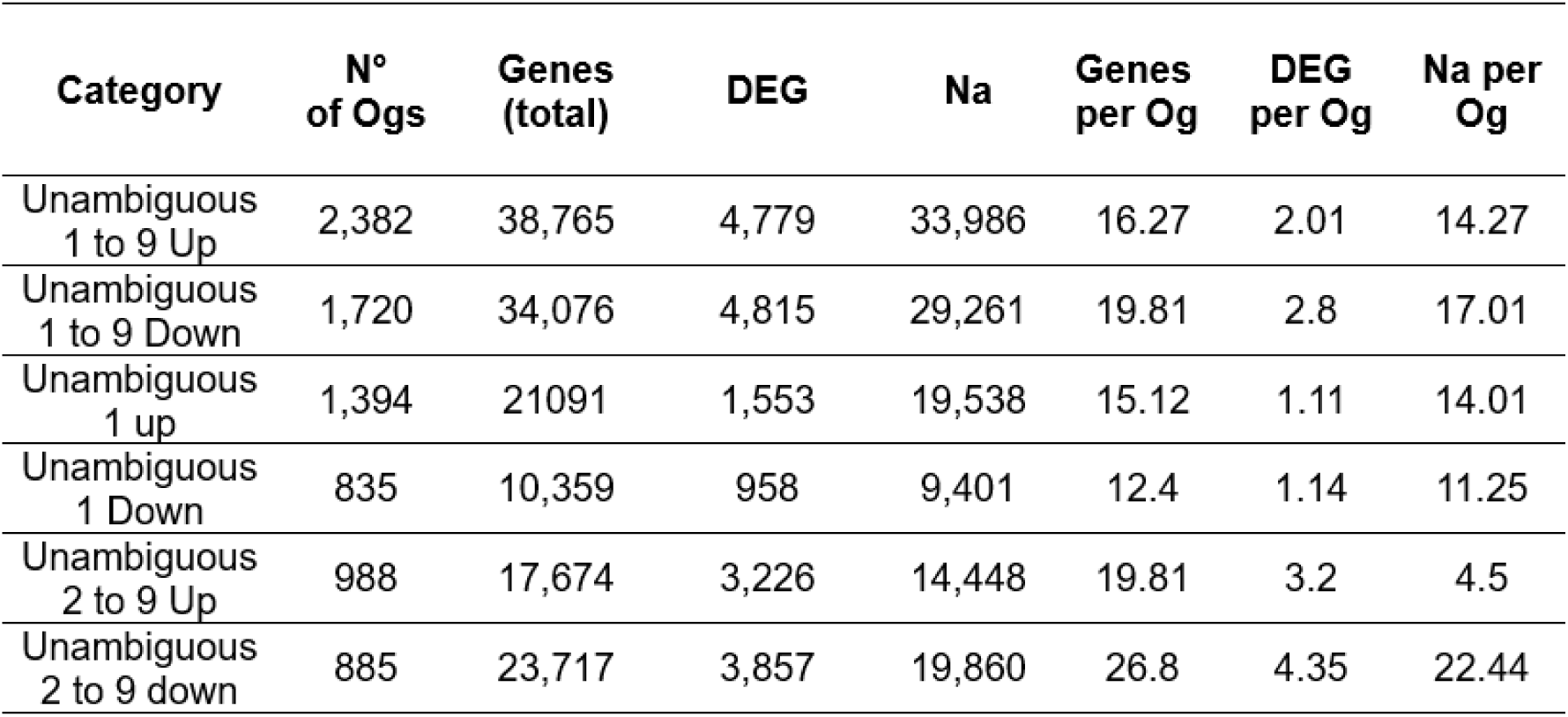
Number of orthogroups (OGs) with unambiguous upregulated or downregulated gene expression, along with the corresponding total number of genes and DEGs.

**Table S3.**
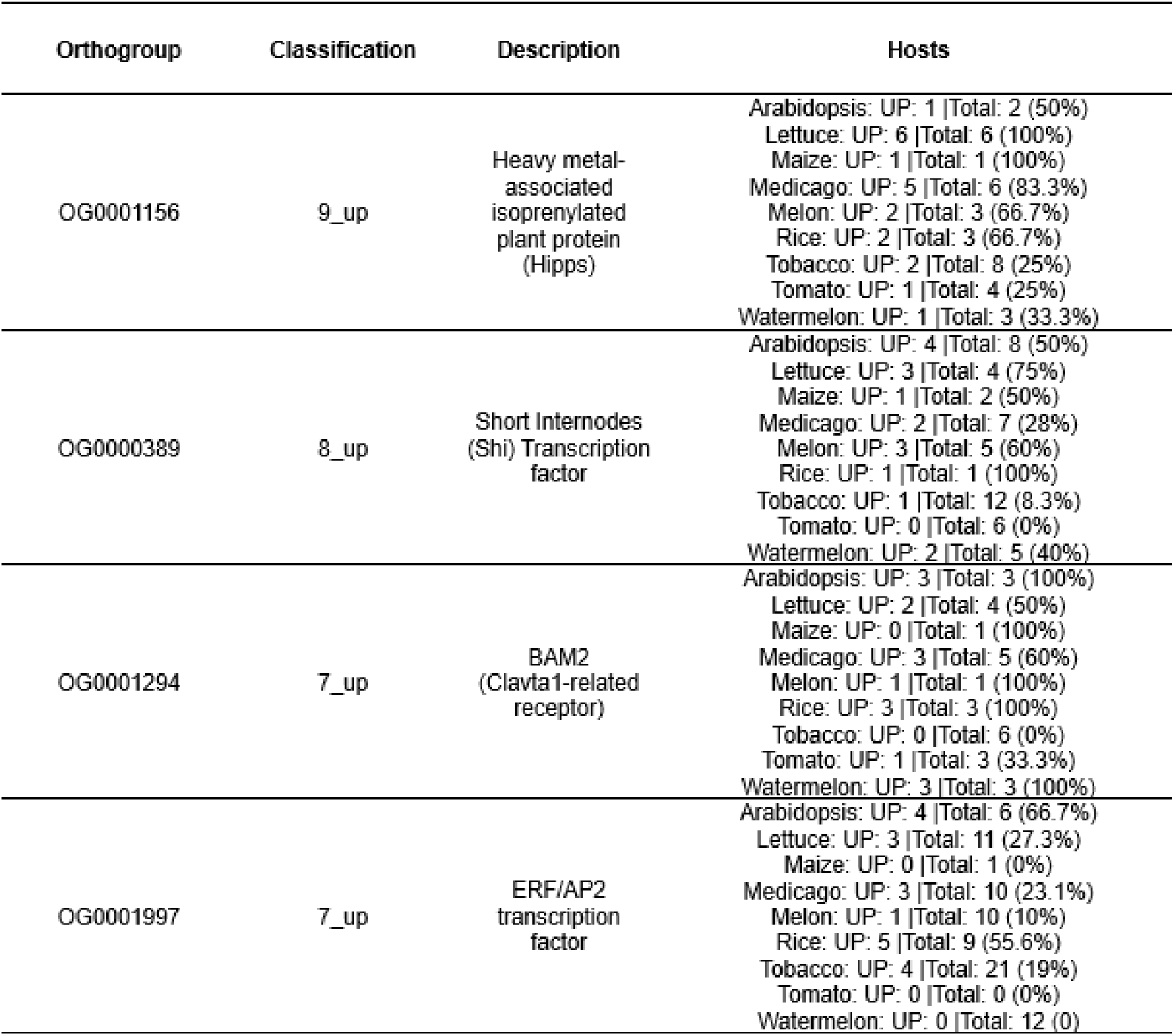
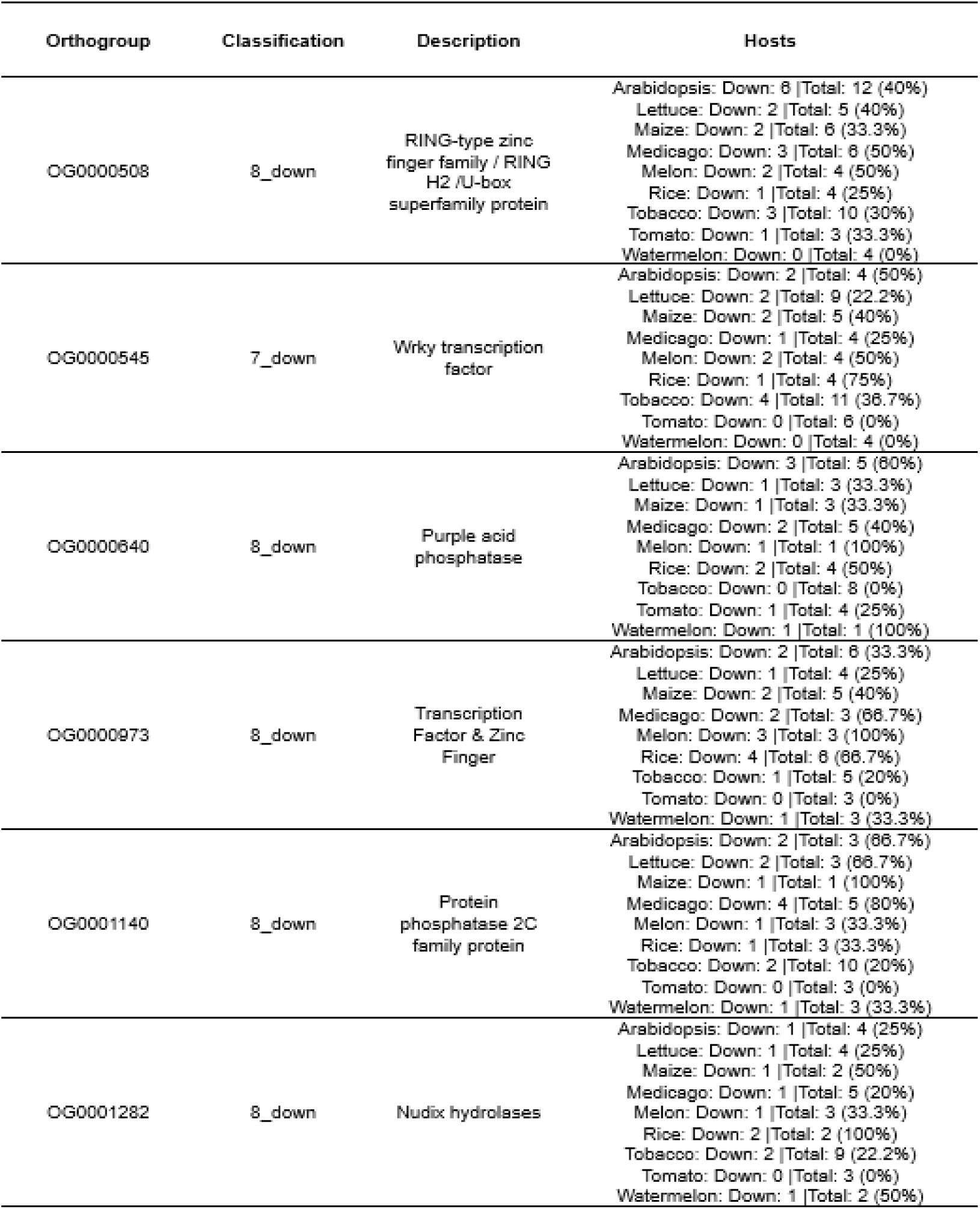
Orthogroups presenting differentially expressed genes, including functional descriptions and gene expression proportions. These orthogroup are shared across all host species. Total refers to the number of genes from each species within a given orthogroup. UP and DOWN indicate the number of genes that were differentially expressed - upregulated or downregulated, respectively.

**Table S4.**
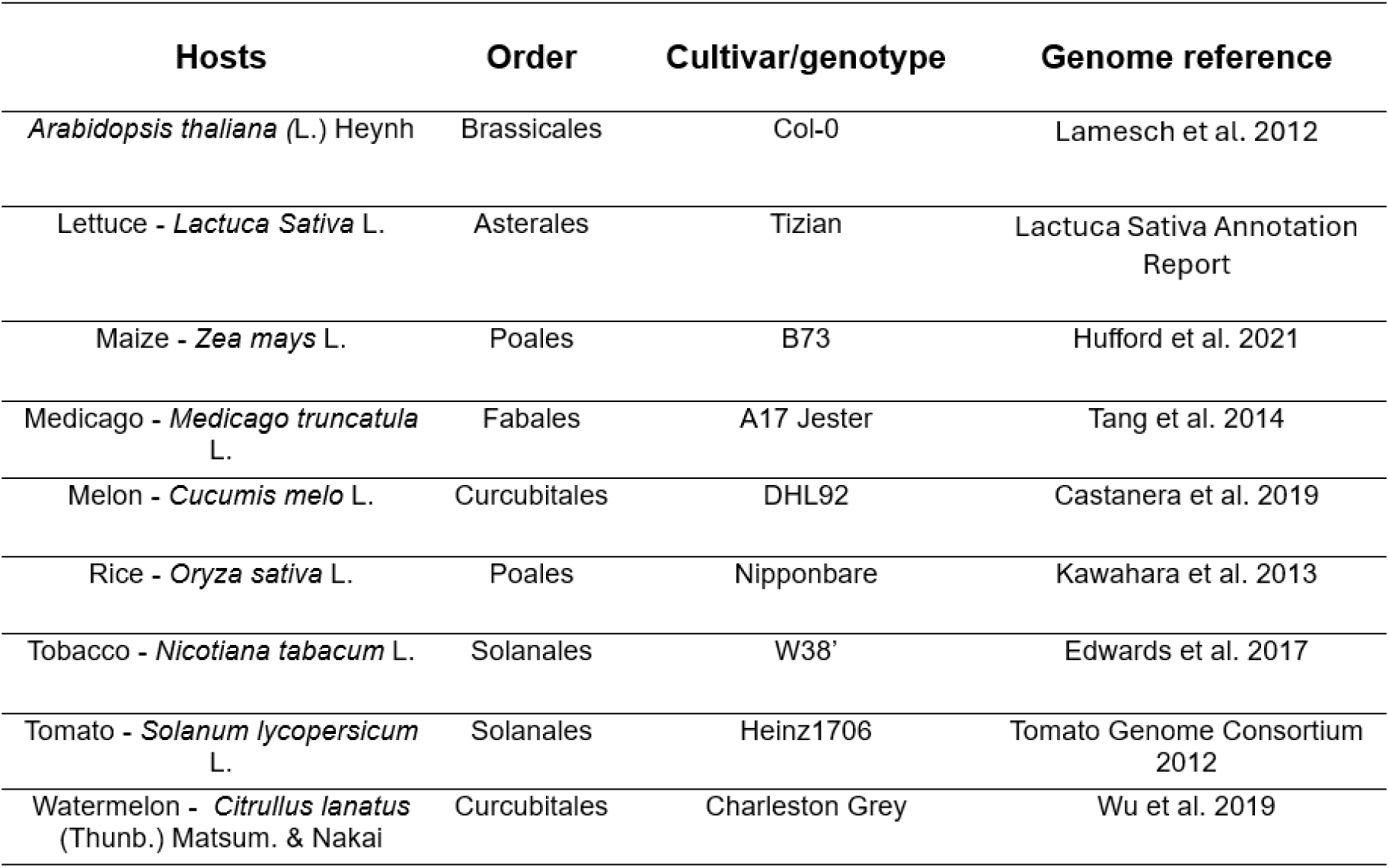
Hosts used in this work, their order, cultivar/genotype and genome reference.

**Table S5.**
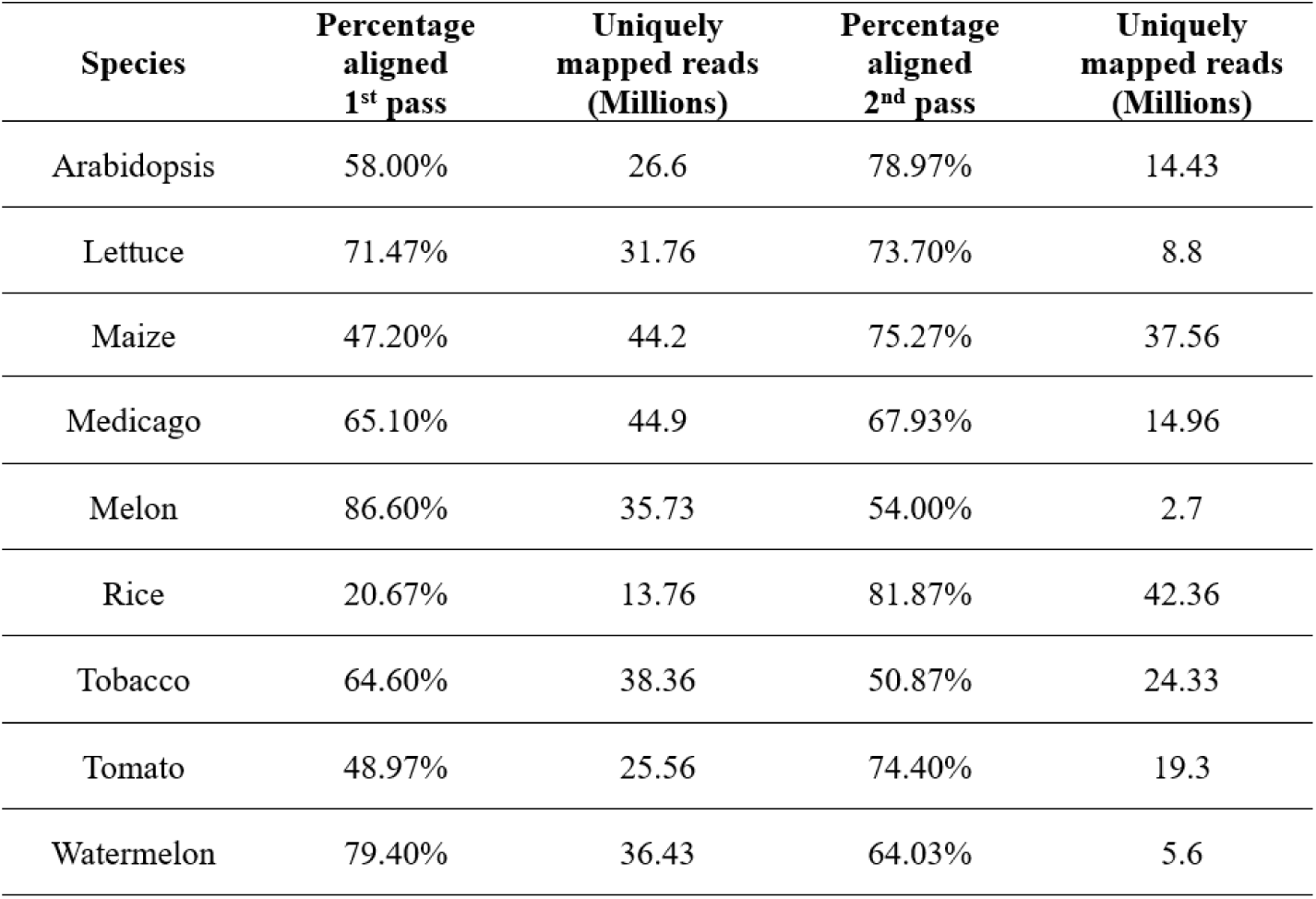
Percentage of mapping and uniquely mapped reads (millions) obtained from multimapping.

## References

1. Gamalero, E. & Glick, B. R. The use of plant growth-promoting bacteria to prevent nematode damage to plants. Biology (Basel) 9, 381 (2020).

2. Eves-van den Akker, S. & Jones, J. T. Sex: Not all that it’s cracked up to be? PLoS Genet. 14, e1007160 (2018).

3. Sacristán, S., Goss, E. M. & den Akker, S. E. How Do Pathogens Evolve Novel Virulence Activities? Molecular Plant-Microbe Interactions (2021) doi:10.1094/MPMI-09-20-0258-IA.

4. Jones, J. T. et al. Top 10 plant-parasitic nematodes in molecular plant pathology. Mol. Plant Pathol. 14, 946–961 (2013).

5. Molloy, B., Baum, T. & den Akker, S. E. Unlocking the development-and physiology-altering ‘effector toolbox’of plant-parasitic nematodes. 39, 732–738 (2023).

6. Tyler, J. Reproduction Without Males in Aseptic Root Cultures of the Root-Knot Nematode. (1933).

7. Lehman, P. S. Galls on aboveground plant parts caused by root-knot nematodes. Nematology Circular vol. 125 2 (1985).

8. Montarry, J. et al. Recent Advances in Population Genomics of Plant-Parasitic Nematodes. Phytopathology 111, 40–48 (2021).

9. Koutsovoulos, G. D. et al. Population genomics supports clonal reproduction and multiple independent gains and losses of parasitic abilities in the most devastating nematode pest. Evol Appl 13, 442–457 (2020).

10. Lunt, D. H., Kumar, S., Koutsovoulos, G. & Blaxter, M. L. The complex hybrid origins of the root knot nematodes revealed through comparative genomics. PeerJ 2, e356 (2014).

11. Blanc-Mathieu, R. et al. Hybridization and polyploidy enable genomic plasticity without sex in the most devastating plant-parasitic nematodes. PLoS Genet 13, e1006777 (2017).

12. Taylor, A. L. & Sasser, J. N. Biology, Identification and Control of Root-Knot Nematodes. (North Carolina State University Graphics, RALEIGH, 1978).

13. Kumar, S., Stecher, G., Suleski, M. & Hedges, S. B. TimeTree: A resource for timelines, timetrees, and divergence times. Mol. Biol. Evol. 34, 1812–1819 (2017).

14. Blondel, V. D., Guillaume, J.-L., Lambiotte, R. & Lefebvre, E. Fast unfolding of communities in large networks. J. Stat. Mech: Theory Exp. P10008 (2008).

15. Nguyen, C.-N. et al. A root-knot nematode small glycine and cysteine-rich secreted effector, MiSGCR1, is involved in plant parasitism. New Phytol 217, 687–699 (2018).

16. Silva, A. C. et al. The DNA methylation landscape of the root-knot nematode-induced pseudo-organ, the gall, in Arabidopsis, is dynamic, contrasting over time, and critically important for successful parasitism. New Phytol 236, 1888–1907 (2022).

17. Dutta, T. K. et al. Functional analysis of a susceptibility gene (HIPP27) in the Arabidopsis thaliana-Meloidogyne incognita pathosystem by using a genome editing strategy. BMC Plant Biology 23, 1–14 (2023).

18. Da Rocha, M. et al. Genome Expression Dynamics Reveal the Parasitism Regulatory Landscape of the Root-Knot Nematode Meloidogyne incognita and a Promoter Motif Associated with Effector Genes. Genes 12, (2021).

19. Dai, D. et al. Unzipped chromosome-level genomes reveal allopolyploid nematode origin pattern as unreduced gamete hybridization. Nat Commun 14, 7156 (2023).

20. Mota, A. P. Z. et al. Unzipped genome assemblies of polyploid root-knot nematodes reveal unusual and clade-specific telomeric repeats. Nat Commun 15, 773 (2024).

21. Siddique, S. et al. The genome and lifestage-specific transcriptomes of a plant-parasitic nematode and its host reveal susceptibility genes involved in trans-kingdom synthesis of vitamin B5. Nat. Commun. 13, 6190 (2022).

22. Andrews, S., Biggins, L., Inglesfield, S., Carr, H. & Montgomery, J. FastQC: A quality control tool for high throughput sequence data. http://www.bioinformatics.babraham.ac.uk/projects/fastqc/ (2010).

23. Kozlowski, D. K. L. et al. Movements of transposable elements contribute to the genomic plasticity and species diversification in an asexually reproducing nematode pest. Evolutionary Applications 14, 1844–1866 (2021).

24. Dobin, A. et al. STAR: ultrafast universal RNA-seq aligner. Bioinformatics 29, 15–21 (2012).

25. Li, B. & Dewey, C. N. RSEM: accurate transcript quantification from RNA-Seq data with or without a reference genome. BMC Bioinformatics 12, 1–16 (2011).

26. Leng, N. et al. EBSeq: an empirical Bayes hierarchical model for inference in RNA-seq experiments. Bioinformatics 29, 1035–1043 (2013).

27. R Core Team. *R: Language Environment Statistical Computing. R Foundation Statistical Computing*. (Vienna, Austria, 2022).

28. Love, M. I., Huber, W. & Anders, S. Moderated estimation of fold change and dispersion for RNA-seq data with DESeq2. Genome Biol 15, 550 (2014).

29. R Core Team. The R Stats Package.

30. Bybd, D. W., Kirkpatrick, T. & Barker, K. R. An improved technique for clearing and staining plant tissues for detection of nematodes. J Nematol 15, 142–143 (1983).

31. Silva, L. J. CRAN: Package laercio. (2010).

32. Dinno, A. Dunn.Test: Dunn’s test of multiple comparisons using rank sums. CRAN: Contributed Packages The R Foundation 10.32614/cran.package.dunn.test (2014).

33. Petersen, T. N., Brunak, S., von Heijne, G. & Nielsen, H. SignalP 4.0: discriminating signal peptides from transmembrane regions. Nature Methods 8, 785–786 (2011).

34. Nielsen, H. Predicting Secretory Proteins with SignalP. Methods Mol Biol 1611, 59–73 (2017).

35. Bernhofer, M. & Rost, B. TMbed: transmembrane proteins predicted through language model embeddings. BMC Bioinformatics 23, 1–19 (2022).

36. Calia, G. et al. Identification and characterization of specific motifs in effector proteins of plant parasites using MOnSTER. *Commun*. Biol. 7, 850 (2024).

37. Camacho, C. et al. BLAST+: architecture and applications. BMC Bioinformatics 10, 1–9 (2009).

38. Molloy, B. et al. The origin, deployment, and evolution of a plant-parasitic nematode effectorome. PLoS Pathog 20, e1012395 (2024).

39. Rizzo, M. & Szekely, G. energy: E-Statistics: Multivariate Inference via the Energy of Data. CRAN: Contributed Packages The R Foundation 10.32614/cran.package.energy (2004).

40. Bastian, M., Heymann, S. & Jacomy, M. Gephi: An open source software for exploring and manipulating networks. Proceedings of the International AAAI Conference on Web and Social Media 3, 361–362 (2009).

41. Shen, W.-K. et al. AnimalTFDB 4.0: a comprehensive animal transcription factor database updated with variation and expression annotations. Nucleic Acids Res 51, D39–D45 (2022).

42. Jones, P. et al. InterProScan 5: genome-scale protein function classification. Bioinformatics 30, 1236–1240 (2014).

43. Urwin, P. E., Lilley, C. J. & Atkinson, H. J. Ingestion of double-stranded RNA by preparasitic juvenile cyst nematodes leads to RNA interference. Mol Plant Microbe Interact 15, 747–752 (2002).

