## Supplementary material for "The host range paradox of *Meloidogyne incognita*: a physiological and transcriptomic analysis of nine susceptible interactions across six plant orders": Table S1

| **Species** | **Coding genes** | **Total DEGs** | **%_total** | **up** | **%_up** | **down** | **%_down** |
| --- | --- | --- | --- | --- | --- | --- | --- |
| Arabidopsis | 27628 | 5638 | 20.4 | 2477 | 9.0 | 3161 | 11.4 |
| Lettuce | 36873 | 3292 | 11.9 | 1536 | 5.6 | 1756 | 6.4 |
| Maize | 39756 | 2591 | 9.4 | 883 | 3.2 | 1708 | 6.2 |
| Melon | 28299 | 2052 | 7.4 | 943 | 3.4 | 1109 | 4.0 |
| Medicago | 50444 | 5063 | 18.3 | 2263 | 8.2 | 2800 | 10.1 |
| Rice | 44844 | 5439 | 19.7 | 2874 | 10.4 | 2565 | 9.3 |
| Tobacco | 69500 | 3780 | 13.7 | 1287 | 4.7 | 2493 | 9.0 |
| Tomato | 34429 | 1398 | 5.1 | 805 | 2.9 | 593 | 2.1 |
| Watermelon | 22764 | 548 | 2.0 | 346 | 1.3 | 202 | 0.7 |

**Table S1.** Total number of coding genes for each species, the number of differentially expressed genes (DEGs), counts of upregulated and downregulated genes, and their respective proportions.
