## Supplementary material for "The host range paradox of *Meloidogyne incognita*: a physiological and transcriptomic analysis of nine susceptible interactions across six plant orders": Table S2

| **Category** | **N°**  **of Ogs** | **Genes (total)** | **DEG** | **Na** | **Genes per Og** | **DEG**  **per Og** | **Na per Og** |
| --- | --- | --- | --- | --- | --- | --- | --- |
| Unambiguous 1 to 9 Up | 2,382 | 38,765 | 4,779 | 33,986 | 16.27 | 2.01 | 14.27 |
| Unambiguous 1 to 9 Down | 1,720 | 34,076 | 4,815 | 29,261 | 19.81 | 2.8 | 17.01 |
| Unambiguous 1 up | 1,394 | 21091 | 1,553 | 19,538 | 15.12 | 1.11 | 14.01 |
| Unambiguous 1 Down | 835 | 10,359 | 958 | 9,401 | 12.4 | 1.14 | 11.25 |
| Unambiguous 2 to 9 Up | 988 | 17,674 | 3,226 | 14,448 | 19.81 | 3.2 | 4.5 |
| Unambiguous 2 to 9 down | 885 | 23,717 | 3,857 | 19,860 | 26.8 | 4.35 | 22.44 |

**Table S2.** Number of orthogroups (OGs) with unambiguous upregulated or downregulated gene expression, along with the corresponding total number of genes and DEGs.
