## Supplementary material for "The host range paradox of *Meloidogyne incognita*: a physiological and transcriptomic analysis of nine susceptible interactions across six plant orders": Table S3

**Table S3.** Orthogroups presenting differentially expressed genes, including functional descriptions and gene expression proportions. These orthogroup are shared across all host species. Total refers to the number of genes from each species within a given orthogroup. UP and DOWN indicate the number of genes that were differentially expressed - upregulated or downregulated, respectively.

Unambiguous up

| **Orthogroup** | **Classification** | **Description** | **Hosts** |
| --- | --- | --- | --- |
| OG0001156 | 9_up | Heavy metal-associated isoprenylated plant protein (Hipps) | Arabidopsis: UP: 1 \|Total: 2 (50%)  Lettuce: UP: 6 \|Total: 6 (100%)  Maize: UP: 1 \|Total: 1 (100%)  Medicago: UP: 5 \|Total: 6 (83.3%)  Melon: UP: 2 \|Total: 3 (66.7%)  Rice: UP: 2 \|Total: 3 (66.7%)  Tobacco: UP: 2 \|Total: 8 (25%)  Tomato: UP: 1 \|Total: 4 (25%)  Watermelon: UP: 1 \|Total: 3 (33.3%) |
| OG0000389 | 8_up | Short Internodes (Shi) Transcription factor | Arabidopsis: UP: 4 \|Total: 8 (50%)  Lettuce: UP: 3 \|Total: 4 (75%)  Maize: UP: 1 \|Total: 2 (50%)  Medicago: UP: 2 \|Total: 7 (28%)  Melon: UP: 3 \|Total: 5 (60%)  Rice: UP: 1 \|Total: 1 (100%)  Tobacco: UP: 1 \|Total: 12 (8.3%)  Tomato: UP: 0 \|Total: 6 (0%)  Watermelon: UP: 2 \|Total: 5 (40%) |
| OG0001294 | 7_up | BAM2  (Clavta1-related receptor) | Arabidopsis: UP: 3 \|Total: 3 (100%)  Lettuce: UP: 2 \|Total: 4 (50%)  Maize: UP: 0 \|Total: 1 (100%)  Medicago: UP: 3 \|Total: 5 (60%)  Melon: UP: 1 \|Total: 1 (100%)  Rice: UP: 3 \|Total: 3 (100%)  Tobacco: UP: 0 \|Total: 6 (0%)  Tomato: UP: 1 \|Total: 3 (33.3%)  Watermelon: UP: 3 \|Total: 3 (100%) |
| OG0001997 | 7_up | ERF/AP2 transcription factor | Arabidopsis: UP: 4 \|Total: 6 (66.7%)  Lettuce: UP: 3 \|Total: 11 (27.3%)  Maize: UP: 0 \|Total: 1 (0%)  Medicago: UP: 3 \|Total: 10 (23.1%)  Melon: UP: 1 \|Total: 10 (10%)  Rice: UP: 5 \|Total: 9 (55.6%)  Tobacco: UP: 4 \|Total: 21 (19%)  Tomato: UP: 0 \|Total: 0 (0%)  Watermelon: UP: 0 \|Total: 12 (0) |

Unambiguous down

| **Orthogroup** | **Classification** | **Description** | **Hosts** |
| --- | --- | --- | --- |
| OG0000508 | 8_down | RING-type zinc finger family / RING H2 /U-box superfamily protein | Arabidopsis: Down: 6 \|Total: 12 (40%)  Lettuce: Down: 2 \|Total: 5 (40%)  Maize: Down: 2 \|Total: 6 (33.3%)  Medicago: Down: 3 \|Total: 6 (50%)  Melon: Down: 2 \|Total: 4 (50%)  Rice: Down: 1 \|Total: 4 (25%)  Tobacco: Down: 3 \|Total: 10 (30%)  Tomato: Down: 1 \|Total: 3 (33.3%)  Watermelon: Down: 0 \|Total: 4 (0%) |
| OG0000545 | 7_down | Wrky transcription factor | Arabidopsis: Down: 2 \|Total: 4 (50%)  Lettuce: Down: 2 \|Total: 9 (22.2%)  Maize: Down: 2 \|Total: 5 (40%)  Medicago: Down: 1 \|Total: 4 (25%)  Melon: Down: 2 \|Total: 4 (50%)  Rice: Down: 1 \|Total: 4 (75%)  Tobacco: Down: 4 \|Total: 11 (36.7%)  Tomato: Down: 0 \|Total: 6 (0%)  Watermelon: Down: 0 \|Total: 4 (0%) |
| OG0000640 | 8_down | Purple acid phosphatase | Arabidopsis: Down: 3 \|Total: 5 (60%)  Lettuce: Down: 1 \|Total: 3 (33.3%)  Maize: Down: 1 \|Total: 3 (33.3%)  Medicago: Down: 2 \|Total: 5 (40%)  Melon: Down: 1 \|Total: 1 (100%)  Rice: Down: 2 \|Total: 4 (50%)  Tobacco: Down: 0 \|Total: 8 (0%)  Tomato: Down: 1 \|Total: 4 (25%)  Watermelon: Down: 1 \|Total: 1 (100%) |
| OG0000973 | 8_down | Transcription Factor & Zinc Finger | Arabidopsis: Down: 2 \|Total: 6 (33.3%)  Lettuce: Down: 1 \|Total: 4 (25%)  Maize: Down: 2 \|Total: 5 (40%)  Medicago: Down: 2 \|Total: 3 (66.7%)  Melon: Down: 3 \|Total: 3 (100%)  Rice: Down: 4 \|Total: 6 (66.7%)  Tobacco: Down: 1 \|Total: 5 (20%)  Tomato: Down: 0 \|Total: 3 (0%)  Watermelon: Down: 1 \|Total: 3 (33.3%) |
| OG0001140 | 8_down | Protein phosphatase 2C family protein | Arabidopsis: Down: 2 \|Total: 3 (66.7%)  Lettuce: Down: 2 \|Total: 3 (66.7%)  Maize: Down: 1 \|Total: 1 (100%)  Medicago: Down: 4 \|Total: 5 (80%)  Melon: Down: 1 \|Total: 3 (33.3%)  Rice: Down: 1 \|Total: 3 (33.3%)  Tobacco: Down: 2 \|Total: 10 (20%)  Tomato: Down: 0 \|Total: 3 (0%)  Watermelon: Down: 1 \|Total: 3 (33.3%) |
| OG0001282 | 8_down | Nudix hydrolases | Arabidopsis: Down: 1 \|Total: 4 (25%)  Lettuce: Down: 1 \|Total: 4 (25%)  Maize: Down: 1 \|Total: 2 (50%)  Medicago: Down: 1 \|Total: 5 (20%)  Melon: Down: 1 \|Total: 3 (33.3%)  Rice: Down: 2 \|Total: 2 (100%)  Tobacco: Down: 2 \|Total: 9 (22.2%)  Tomato: Down: 0 \|Total: 3 (0%)  Watermelon: Down: 1 \|Total: 2 (50%) |
| OG0001943 | 8_down | ERF/AP2 transcription factor | Arabidopsis: Down: 1 \|Total: 5 (20%)  Lettuce: Down: 2 \|Total: 3 (66.7%)  Maize: Down: 2 \|Total: 5 (40%)  Medicago: Down: 1 \|Total: 3 (33.3%)  Melon: Down: 1 \|Total: 3 (33.3%)  Rice: Down: 1 \|Total: 2 (50%)  Tobacco: Down: 2 \|Total: 6 (33.3%)  Tomato: Down: 1 \|Total: 3 (33.3%)  Watermelon: Down: 0 \|Total: 3 (0%) |
| OG0002374 | 8_down | MYB transcription factor | Arabidopsis: Down: 2 \|Total: 3 (66.7%)  Lettuce: Down: 1 \|Total: 2 (50%)  Maize: Down: 1 \|Total: 1 (100%)  Medicago: Down: 1 \|Total: 4 (25%)  Melon: Down: 1 \|Total: 3 (33.3%)  Rice: Down: 1 \|Total: 3 (33.3%)  Tobacco: Down: 2 \|Total: 6 (33.3%)  Tomato: Down: 1 \|Total: 2 (50%)  Watermelon: Down: 0 \|Total: 3 (0%) |
| OG0002769 | 8_down | E3 ubiquitin transferase  Pub20/CMPG1 activation of defence mechanisms | Arabidopsis: Down: 2 \|Total: 2 (100%)  Lettuce: Down: 2 \|Total: 2 (100%)  Maize: Down: 1 \|Total: 3 (33.3%)  Medicago: Down: 3 \|Total: 4 (75%)  Melon: Down: 2 \|Total: 2 (100%)  Rice: Down: 1 \|Total: 4 (25%)  Tobacco: Down: 4 \|Total: 6 (66.7%)  Tomato: Down: 1 \|Total: 3 (33.3%)  Watermelon: Down: 0 \|Total: 2 (0%) |
| OG0003161 | 8_down | P-Loop containing nucleoside triphosphate hydrolases | Arabidopsis: Down: 1 \|Total: 1 (100%)  Lettuce: Down: 1 \|Total: 2 (50%)  Maize: Down: 1 \|Total: 2 (50%)  Medicago: Down: 3 \|Total: 3 (100%)  Melon: Down: 1 \|Total: 2 (50%)  Rice: Down: 1 \|Total: 3 (33.3%)  Tobacco: Down: 1 \|Total: 6 (16.7%)  Tomato: Down: 2 \|Total: 3 (66.7%)  Watermelon: Down: 0 \|Total: 2 (0%) |
| OG0005554 | 8_down | Glycosyltransferase family | Arabidopsis: Down: 2 \|Total: 3 (66.7%)  Lettuce: Down: 1 \|Total: 2 (50%)  Maize: Down: 1 \|Total: 1 (66.7%)  Medicago: Down: 1 \|Total: 1 (100%)  Melon: Down: 1 \|Total: 1 (100%)  Rice: Down: 1 \|Total: 1 (100%)  Tobacco: Down: 1 \|Total: 4 (25%)  Tomato: Down: 0 \|Total: 1 (0%)  Watermelon: Down: 1 \|Total: 1 (100%) |
| OG0005829 | 8_down | Aspartyl protease family protein | Arabidopsis: Down: 2 \|Total: 2 (100%)  Lettuce: Down: 3 \|Total: 3 (100%)  Maize: Down: 2 \|Total: 3 (66.7%)  Medicago: Down: 2 \|Total: 3 (66.7%)  Melon: Down: 1 \|Total: 1 (100%)  Rice: Down: 1 \|Total: 1 (100%)  Tobacco: Down: 2 \|Total: 4 (50%)  Tomato: Down: 1 \|Total: 2 (50%)  Watermelon: Down: 0 \|Total: 1 (0%) |
| OG0008161 | 8_down | Thylakoidal processing peptidase | Arabidopsis: Down: 1 \|Total: 2 (50%)  Lettuce: Down: 1 \|Total: 2 (50%)  Maize: Down: 1 \|Total: 1 (100%)  Medicago: Down: 2 \|Total: 3 (66.7%)  Melon: Down: 1 \|Total: 2 (50%)  Rice: Down: 1 \|Total: 1 (100%)  Tobacco: Down: 1 \|Total: 2 (50%)  Tomato: Down: 1 \|Total: 1 (100%)  Watermelon: Down: 0 \|Total: 1 (0%) |
| OG0010480 | 8_down | Small Auxin UP RNA (SAUR) E3 ubiquitin ligase Plant U box | Arabidopsis: Down: 1 \|Total: 1 (100%)  Lettuce: Down: 1 \|Total: 1 (100%)  Maize: Down: 2 \|Total: 3 (66.7%)  Medicago: Down: 1 \|Total: 1 (100%)  Melon: Down: 1 \|Total: 1 (100%)  Rice: Down: 2 \|Total: 2 (100%)  Tobacco: Down: 1 \|Total: 2 (50%)  Tomato: Down: 1 \|Total: 1 (100%)  Watermelon: Down: 0 \|Total: 1 (0%) |
