## Supplementary material for "The host range paradox of *Meloidogyne incognita*: a physiological and transcriptomic analysis of nine susceptible interactions across six plant orders": Table S4

**Table S4.** Hosts used in this work, their order, cultivar/genotype and genome reference.

| **Hosts** | **Order** | **Cultivar/genotype** | **Genome reference** |
| --- | --- | --- | --- |
| *Arabidopsis thaliana (*L.) Heynh | Brassicales | Col-0 | Lamesch et al. 2012 |
| Lettuce - *Lactuca Sativa* L. | Asterales | Tizian | Lactuca Sativa Annotation Report |
| Maize - *Zea mays* L. | Poales | B73 | Hufford et al. 2021 |
| Medicago - *Medicago truncatula* L. | Fabales | A17 Jester | Tang et al. 2014 |
| Melon - *Cucumis melo* L. | Curcubitales | DHL92 | Castanera et al. 2019 |
| Rice - *Oryza* *sativa* L. | Poales | Nipponbare | Kawahara et al. 2013 |
| Tobacco - *Nicotiana tabacum* L. | Solanales | W38’ | Edwards et al. 2017 |
| Tomato - *Solanum lycopersicum* L. | Solanales | Heinz1706 | Tomato Genome Consortium 2012 |
| Watermelon -  *Citrullus lanatus* (Thunb.) Matsum. & Nakai | Curcubitales | Charleston Grey | Wu et al. 2019 |
