## Supplementary material for "The host range paradox of *Meloidogyne incognita*: a physiological and transcriptomic analysis of nine susceptible interactions across six plant orders": Table S5

| **Species** | **Percentage aligned 1^st^ pass** | **Uniquely mapped reads (Millions)** | **Percentage aligned 2^nd^ pass** | **Uniquely mapped reads (Millions)** |
| --- | --- | --- | --- | --- |
| Arabidopsis | 58.00% | 26.6 | 78.97% | 14.43 |
| Lettuce | 71.47% | 31.76 | 73.70% | 8.8 |
| Maize | 47.20% | 44.2 | 75.27% | 37.56 |
| Medicago | 65.10% | 44.9 | 67.93% | 14.96 |
| Melon | 86.60% | 35.73 | 54.00% | 2.7 |
| Rice | 20.67% | 13.76 | 81.87% | 42.36 |
| Tobacco | 64.60% | 38.36 | 50.87% | 24.33 |
| Tomato | 48.97% | 25.56 | 74.40% | 19.3 |
| Watermelon | 79.40% | 36.43 | 64.03% | 5.6 |

**Table S5**. Percentage of mapping and uniquely mapped reads (millions) obtained from multimapping.
